# Epigenetic silencing of MAFG is a potential prognosis biomarker in lung adenocarcinomas

**DOI:** 10.64898/2026.05.26.724922

**Authors:** Alvaro Garcia-Guede, Carlos Rodríguez-Antolín, Ana Arauzo-Cabrera, Rocío Moreno-Velasco, Olga Pernía, Miranda Burdiel, Lucía Acero-Riaguas, Isabel Esteban-Rodríguez, Silvia Sacristán, Raúl Torres-Ruiz, Sandra Rodriguez-Perales, Ana Sastre-Perona, Victor M. González, Javier De Castro, Inmaculada Ibáñez de Cáceres, Olga Vera

## Abstract

Non-small cell lung cancer (NSCLC) remains one of the leading causes of cancer-related mortality, partly because it is often diagnosed at advanced stages and frequently develops resistance to platinum-based chemotherapy. We previously showed that MAFG becomes derepressed following miR-7 hypermethylation, promoting platinum resistance in NSCLC and ovarian cancer cell lines. Although MAFG is a well-established regulator of oxidative stress, recent evidence in melanoma and colorectal cancer suggests an additional role as a regulator of methylator phenotypes. However, how MAFG reshapes the lung cancer epigenome remains unknown. Here, we investigated the contribution of MAFG to DNA methylation remodeling by combining CRISPR/Cas9-mediated MAFG deletion with CpG-Methyl-Array profiling, followed by expression (qPCR) and methylation (qMSP) validation in tumor cell lines. Our translational approach integrated aptahistochemistry using MAFG-specific aptamers in 127 NSCLC patients, methylation analysis in 35 fresh-frozen tumors and 40 FFPE samples, and interrogation of TCGA methylation datasets. MAFG loss reduced promoter methylation of *LIF* and *MAFG* itself. Importantly, these effects were subtype-specific, with MAFG expression and methylation displaying distinct transcriptional programs in LUAD versus LUSC, and prognostic associations restricted to KRAS-mutated adenocarcinomas. In NSCL in silico and in house cohorts, lower MAFG methylation and higher MAFG protein levels were both associated with worse prognosis. In summary, our findings identify MAFG as a regulator of DNA methylation in NSCLC and support the use of MAFG DNA methylation, or protein levels as clinically relevant prognostic biomarkers, particularly in lung adenocarcinoma.

## INTRODUCTION

Lung cancer remains the leading cause of cancer-related mortality worldwide, accounting for over one million deaths annually worldwide^1^. Non-small cell lung cancer (NSCLC) represents more than 80% of lung cancer cases and is primarily classified into lung adenocarcinoma (LUAD) and lung squamous cell carcinoma (LUSC). Despite therapeutic advances, platinum-based chemotherapy continues to be the cornerstone of treatment. However, continued exposure to these agents promotes adaptive resistance mechanisms, and the disease almost invariably progresses to a platinum-resistant state. Thus, understanding the molecular mechanisms of resistance is essential for developing more effective therapeutic strategies and identifying diagnostic and prognostic biomarkers.

In recent years, epigenetic regulation has emerged as a critical layer of control in cell biology and disease development^2,3^. While genetic alterations are well-characterized in early tumorigenesis, non-genetic alterations that drive progression to advanced stages remain poorly understood. Epigenetic mechanisms such as DNA methylation and histone posttranslational modifications, which promote chromatin condensation, along with posttranscriptional regulation by non-coding RNAs, have traditionally been recognized as key modulators of gene expression^2,3^. Advances in high-throughput technologies have recently uncover a variety of epifactors that influence transcriptional activity without altering the genomic sequence^4^.

In our search for potential epifactors influencing chemotherapy resistance in cancer, we previously identified the small MAF transcription factor, MAFG, as a target of microRNA-7 in cisplatin-sensitive lung and ovarian cancer cell lines^5,6^. Continuous exposure to cisplatin induces the epigenetic silencing of miR-7, leading to MAFG de-repression and contributing to the development of resistance against platinum-based therapies by protecting from drug-induced reactive oxygen species (ROS) production^5,6^. Similar findings have been reported in other tumor types^7^. In cancer, MAFG has been identified as a poor prognosis marker in breast and melanoma cancers, where MAFG promotes tumor cell dedifferentiation^8,9^. Furthermore, its regulation by MAT2A contributes to liver damage and hepatocellular carcinoma development^10^. Beyond its role as a transcriptional co-activator of oxidative response genes^11^, MAFG has recently been shown to recruit an epigenetic silencing complex –formed by BACH1, CDH8 and DNMT3B – in colorectal cancer and melanoma, promoting the repression of frequently methylated genes^12,13^. In autoimmune encephalomyelitis models, MAFG also represses genes involved in inflammation and immune regulation^14^. Despite the growing evidence of its involvement in cancer development and epigenetic control, the molecular mechanisms underlying the regulatory functions of MAFG and their impact across oncological contexts remain largely unexplored. In this study, we investigated the epigenetic role of MAFG as a DNA-methylating agent and assessed its clinical and therapeutic relevance in lung cancer.

## RESULTS

### MAFG deletion impacts the DNA methylation landscape of epithelial cancer cells

To investigate the epigenetic role of MAFG in cancer progression and therapy resistance, we established a loss-of-function model using CRISPR/Cas9 in cisplatin-resistant H23R and A2780R cell lines, which exhibit high MAFG expression^5^, and the NSCLC cell line, A549. We designed specific sgRNAs (sg1 and sg2) targeting the MAFG coding to maximize editing efficiency while minimizing off-target activity (**Supplementary Figure 1A**). Deletion screening was performed by PCR using primers F1-R2 on individual clones, spanning the genomic region targeted by the sgRNAs. Successful MAFG knock-out (KO) was confirmed in four H23R clones (**Supplementary Figure 1B**), four A549 clones (**Supplementary Figure 1C**), and seven A2780R clones (**Supplementary Figure 1D**), although only two A2780R clones remained viable for downstream experiments. We validated MAFG downregulation by qRT-PCR in three clones for H23R (**Figure 1A**), and in all A549 (**Figure 1B**) and A2780R clones (**Figure 1C**), using a wild-type clone (Ctrl) that underwent the same CRISPR/Cas9 procedure without effective deletion as a control. One KO clone per cell line was selected for further analyses, and maintenance of the MAFG deletion was confirmed by PCR after culture expansion using primers F1–R2 (**Figure 1D**). Additional PCRs using primer pairs F1–R1 and F2–R2 (**Supplementary Figure 1A** and **1E**), confirmed homozygous KO in H23R and A549, and heterozygous KO in A2780R. Sanger sequencing of the amplified region confirmed the expected deletion within the MAFG locus (**Figure 1E**), and was accompanied by reduced MAFG protein levels (**Figure 1F**).

**Figure 1.**
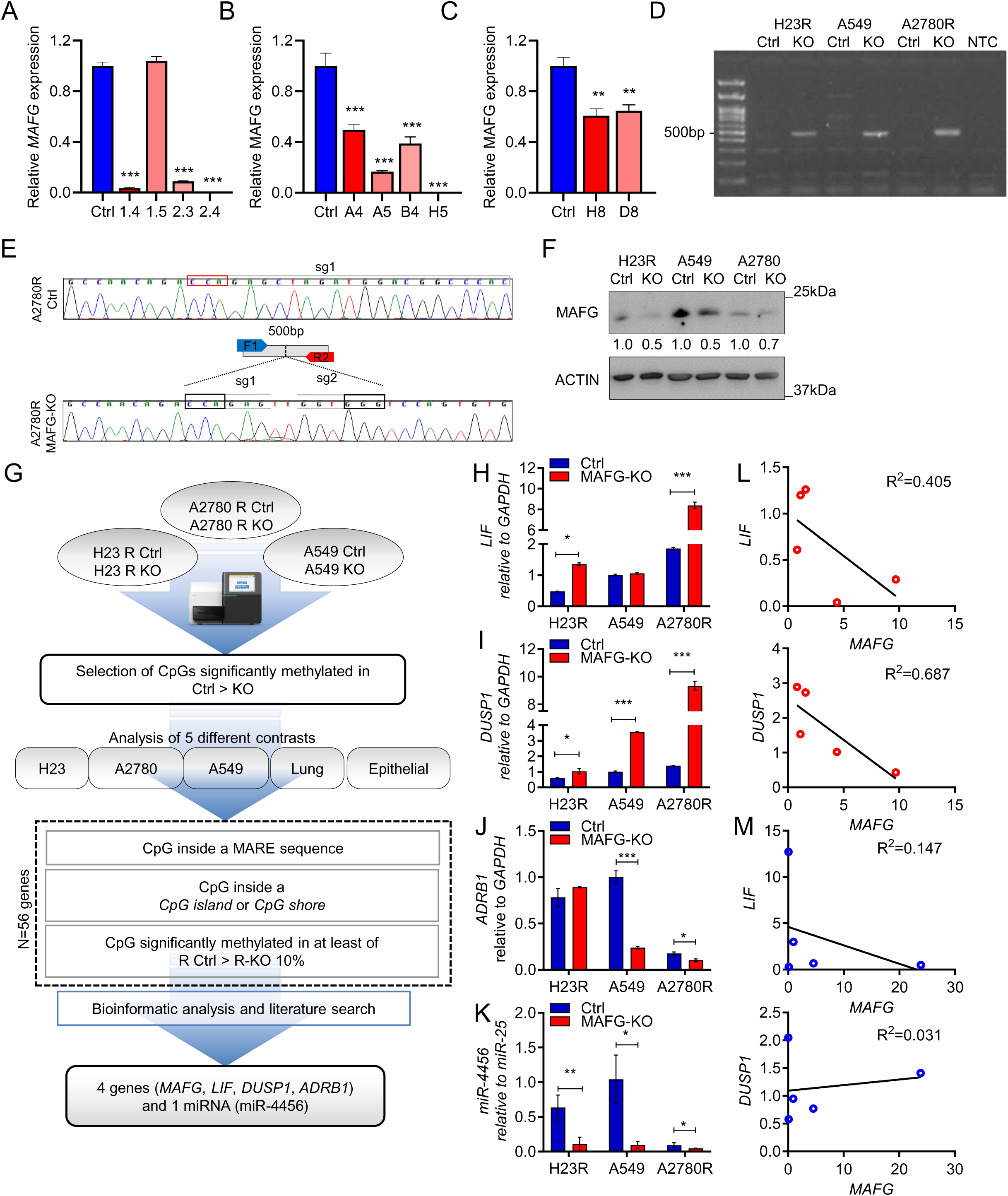
Generation of MAFG knockout models and identification of epigenetically regulated target genes. **(A-C)** qRT-PCR validating KO of *MAFG* in H23R (A), A549 (B) and A2780R (C) cell lines. n = 3 technical replicates from one out of two biological replicates are represented. Statistical significance was determined using Welch’s two-tailed t-test using the Ctrl group as reference. Error bars represent mean + s.d. **p<0.01; ***p<0.001. **(B)** Semi-quantitative PCR amplification using primer pairs F1–R2 in genomic DNA from Ctrl and MAFG-KO clones derived from H23R, A2780R and A549 cell lines. **(C)** Representative Sanger sequencing of PCR products from Ctrl and MAFG-KO clones, confirming deletion of the targeted region. sgRNA target sequences are highlighted with a grey rectangle and PAM sequences in black rectangle. **(D)** Western blot validation of MAFG KO in H23R, A549 and A2780R cell lines. Blot shown represents one out of three experiments with the same result. Numbers represent the ratio of the band intensity between MAFG and ACTIN. **(F)** Workflow describing the bioinformatic strategy used to identify differentially methylated CpG sites and candidate genes from SeqCap methylome profiling in Ctrl and MAFG-KO clones. **(H-K)** qRT-PCR validating the changes in expression in LIF (H), DUSP1 (I), ADRB1 (J) and miR-4456 (K) in Ctrl and MAFG-KO clones from H23R, A2780R and A549 cell lines. For gene expression, *GAPDH* was used as endogenous control, whereas miR-25 was used as housekeeping miRNA in (K). n = 3 technical replicates from one out of two biological replicates are represented. Statistical significance was determined using Welch’s two-tailed t-test using the Ctrl group as reference. Error bars represent mean + s.d. *p<0.05; **p<0.01; ***p<0.001. **(L-M)** Linear regression analysis of *MAFG* expression with *LIF* and *DUSP1* in fresh tumor tissues from lung adenocarcinoma (LUAD, n=5) (L) and lung squamous cell carcinoma (LUSC, n=5) (M) patients. R² indicates the coefficient of determination.

After establishing the MAFG-KO models, we examined the epigenetic consequences of MAFG loss, focusing on DNA methylation. Using SeqCap technology, we assessed the methylation levels at 5.5 million CpG positions across the genome in the wild-type MAFG (Ctrl) cell lines and their MAFG-KO counterparts. Our bioinformatic analysis included: (i) individual comparisons for each cell line, (ii) a combined analysis of the two NSCLC cell lines (H23R and A549), referred to as “Lung”, and (iii) a combined analysis of all three cell lines (H23R, A2780R, and A549), referred to as “Epithelial”, resulting in a total of five datasets (**Figure 1G**). We focused on CpG sites showing significantly decreased methylation in MAFG-KO cells relative to controls, specifically those located within MAF recognition elements (MAREs) in promoter regions, as well as CpG islands or shores of coding or non-coding regions, with a minimum methylation change of 10%. This approach identified 114 CpG positions associated with 56 candidate genes (**Supplementary Table 1**). As pathway enrichment analysis of the 56 candidate genes did not reveal significant associations with known biological pathways (**Supplementary Figure S2A**), we prioritized genes showing differential methylation in at least four of the five analyses and present in both the Lung and Epithelial datasets (**Figure 1G** and **Supplementary Figure S2B**). Additionally, we performed a literature-based prioritization to identify genes with strong links to cancer in general and lung cancer in particular, highlighting *LIF1*, *DUSP1*, *GRHL3*, *MAFG*, *ADRB1* and *WINK2* (**Supplementary Figure S2B,S2C**). Differential methylation of *ADRB1* and microRNA-4456 was observed in the Epithelial, Lung, H23R, and A549 analyses, whereas *LIF* showed differential methylation in the Epithelial, Lung, A549, and A2780R analyses (**Supplementary Figure S2B**). Notably, differential methylation of *MAFG* itself was detected in the Epithelial, Lung, H23R, and A2780R datasets. Based on these results, we focused on *MAFG*, *LIF*, *DUSP1*, *ADRB1*, and microRNA-4456 as the most promising candidates for further investigation.

Initial validation by qRT-PCR in MAFG-KO versus Ctrl cells confirmed increased expression of *LIF* (**Figure 1H**) and *DUSP1* (**Figure 1I**) in H23R and A2780R, whereas *ADRB1* and miR-4456 showed no significant changes (**Figure 1J,K**). As expected, MAFG expression did not increase in KO cells due to the absence of its coding sequence (**Figure 1A-1F**). To evaluate the translational relevance of these findings, we measured *MAFG*, *LIF*, and *DUSP1* expression by qRT-PCR in a pilot cohort of 10 lung cancer patients from University Hospital La Paz (5 lung adenocarcinoma and 5 lung squamous cell carcinoma). In lung adenocarcinoma samples, we observed a negative association between *MAFG* and *LIF* (R² = 0.405), as well as between *MAFG* and *DUSP1* (R² = 0.688) (**Figure 1L**), whereas no association was detected in lung squamous cell carcinoma (**Figure 1M**). These results suggest that MAFG may regulate *LIF* and *DUSP1* specifically in lung adenocarcinoma.

### LIF promoter methylation is associated with MAFG loss in lung cancer cell lines

Following the identification and initial validation of *LIF* and *DUSP1* as candidate genes potentially associated with MAFG-dependent epigenetic regulation, we performed a detailed CpG-level analysis of methylation changes affecting these loci. Using SeqCap data, we examined methylation differences across all CpG positions associated with *LIF* and *DUSP1* in MAFG-KO and control cells (**Figure 2** and **Supplementary Figure S3**). For *LIF*, widespread promoter demethylation was observed in the MAFG-KO groups of A549 and A2780R cells compared with controls, whereas only a single CpG position displayed differential methylation in H23R MAFG-KO cells (**Figure 2A**). In contrast, CpG positions associated with *DUSP1* showed predominantly increased methylation in the absence of MAFG in H23R and A2780R, with no differentially methylated CpGs detected in A549 cells (**Supplementary Figure S3**). Only four positions (three in H23R and one in A2780R) exhibited reduced methylation upon MAFG deletion; however, these were located outside the *DUSP1* promoter region (**Supplementary Figure S3**). Based on the limited number of CpGs and their distal genomic location, *DUSP1* was discarded for further validation.

**Figure 2.**
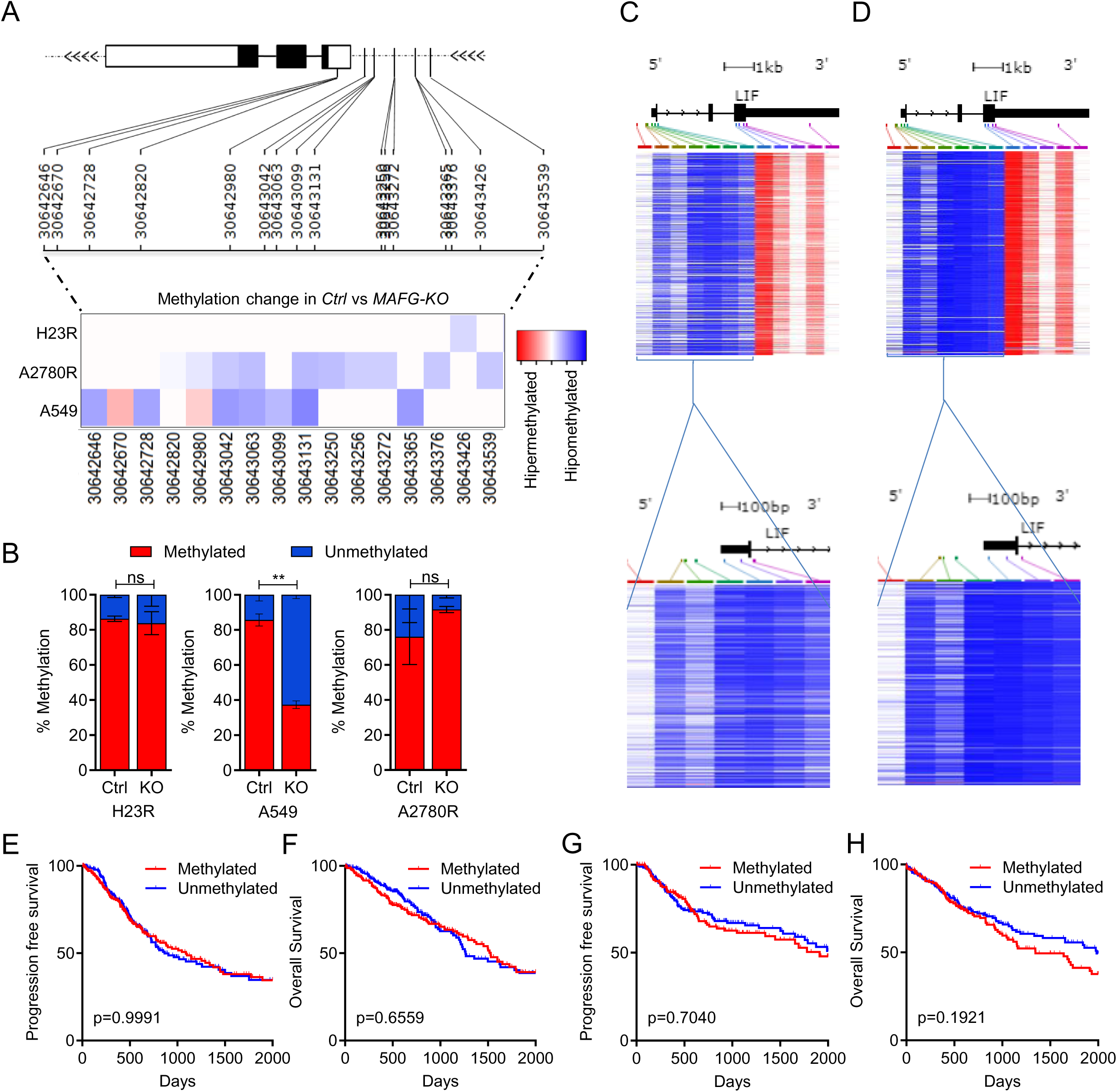
LIF promoter methylation is associated with MAFG loss in lung cancer cell lines. **(A)** Methylation changes observed in SeqCap of CpG sites located in the promoter regions of LIF MAFG-KO vs Ctrl cells. **(B)** Validation of LIF promoter methylation changes by quantitative methylation-specific PCR (qMSP) in H23R, A2780R and A549 cell lines. n = 2 technical replicates from one out of two biological replicates are represented. Statistical significance was determined using Welch’s two-tailed t-test using the Ctrl group as reference. Error bars represent mean + s.d. *p<0.05. **(C,D)** Methylation levels of LIF-associated CpG islands in the TCGA LUAD (C) and LUSC (D) data available at https://xenabrowser.net/. **(E-H)** Kaplan–Meier analysis of progression-free survival (PFS) and overall survival (OS) in the TCGA LUAD (E,F) and LUSC (G,H) data based on methylation levels of CpG sites within the LIF promoter from E and D using https://xenabrowser.net/. Cut-off Beta-value for LUAD is 0.1625 and for LUSC is 0.1173. LIF Methylated in LUAD n = 244; LIF Unmethylated in LUAD n = 239; LIF Methylated in LUSC n = 167; LIF Unmethylated in LUSC n = 198. Log-rank (Mantel-Cox) test was used to determine differences in survival between groups.

To independently validate *LIF* methylation changes, we designed TaqMan probes and primers flanking the CpG positions of interest and assessed DNA methylation levels by quantitative methylation-specific PCR (qMSP). qMSP analyses confirmed a significant reduction in *LIF* promoter methylation (∼60%) in the A549 MAFG-KO cells (**Figure 2B**). In contrast, methylation differences observed in A2780R cells by SeqCap were not validated by qMSP, likely reflecting the smaller magnitude of methylation changes in this cell line (**Figure 2B**). Consistent with the methylome data, no significant methylation changes were detected in H23R MAFG-KO clones (**Figure 2B**). To assess the clinical relevance of *LIF* promoter methylation in NSCLC, we analyzed two TCGA cohorts with available HM450k methylation data: lung adenocarcinoma (LUAD, n = 492) (**Figure 2C**) and lung squamous cell carcinoma (LUSC, n = 415) (**Figure 2D**). CpG positions within the *LIF* promoter region closest to those identified in our experimental models were selected for analysis. Kaplan–Meier survival analyses revealed that *LIF* promoter methylation did not significantly stratify progression-free survival or overall survival in either LUAD (**Figure 2E, 2F**) or LUSC patients (**Figure 2G, 2H**), suggesting that methylation at these specific CpG sites may not have prognostic value in NSCLC.

### MAFG is epigenetically self-regulated and is associated with poor prognosis in lung adenocarcinoma

The identification of MAFG as one of the genes displaying differential methylation following MAFG deletion prompted us to investigate a potential autoregulatory mechanism mediated by DNA methylation. We first examined all CpG positions associated with MAFG that showed significant methylation changes between MAFG-KO and control groups in the SeqCap analysis (**Figure 3A**). In A549 MAFG-KO cells, no sequencing reads were detected in the genomic region flanked by the sgRNAs, consistent with deletion of this locus. In contrast, both H23R and A2780R MAFG-KO cells retained sequencing coverage at the MAFG locus, albeit with markedly fewer reads compared to their respective control counterparts. CpG-level analysis revealed 20 differentially methylated CpG sites in H23R cells and 29 in A2780R cells, with all 20 H23R CpGs shared between the two cell lines (**Figure 3B**). Notably, 15 of these shared CpGs were located within a regulatory region of MAFG, specifically in intron 1, forming a cluster of consecutive CpG sites that exhibited consistent loss of methylation following MAFG deletion. Quantitative analysis of these 15 CpGs showed a 31% decrease in methylation in H23R MAFG-KO cells and a 46% decrease in A2780R MAFG-KO cells relative to their respective controls (**Figure 3B,C**). To validate these results using an independent method, we assessed MAFG methylation by qMSP. This analysis confirmed a pronounced reduction in methylation levels in H23R MAFG-KO cells compared with H23R control cells (**Figure 3D**). Similarly, a 20% reduction in MAFG methylation was observed in A2780R MAFG-KO cells relative to A2780R controls (**Figure 3D**). Together, these results corroborated the SeqCap findings and established a reproducible technique for determining MAFG specific methylation.

**Figure 3.**
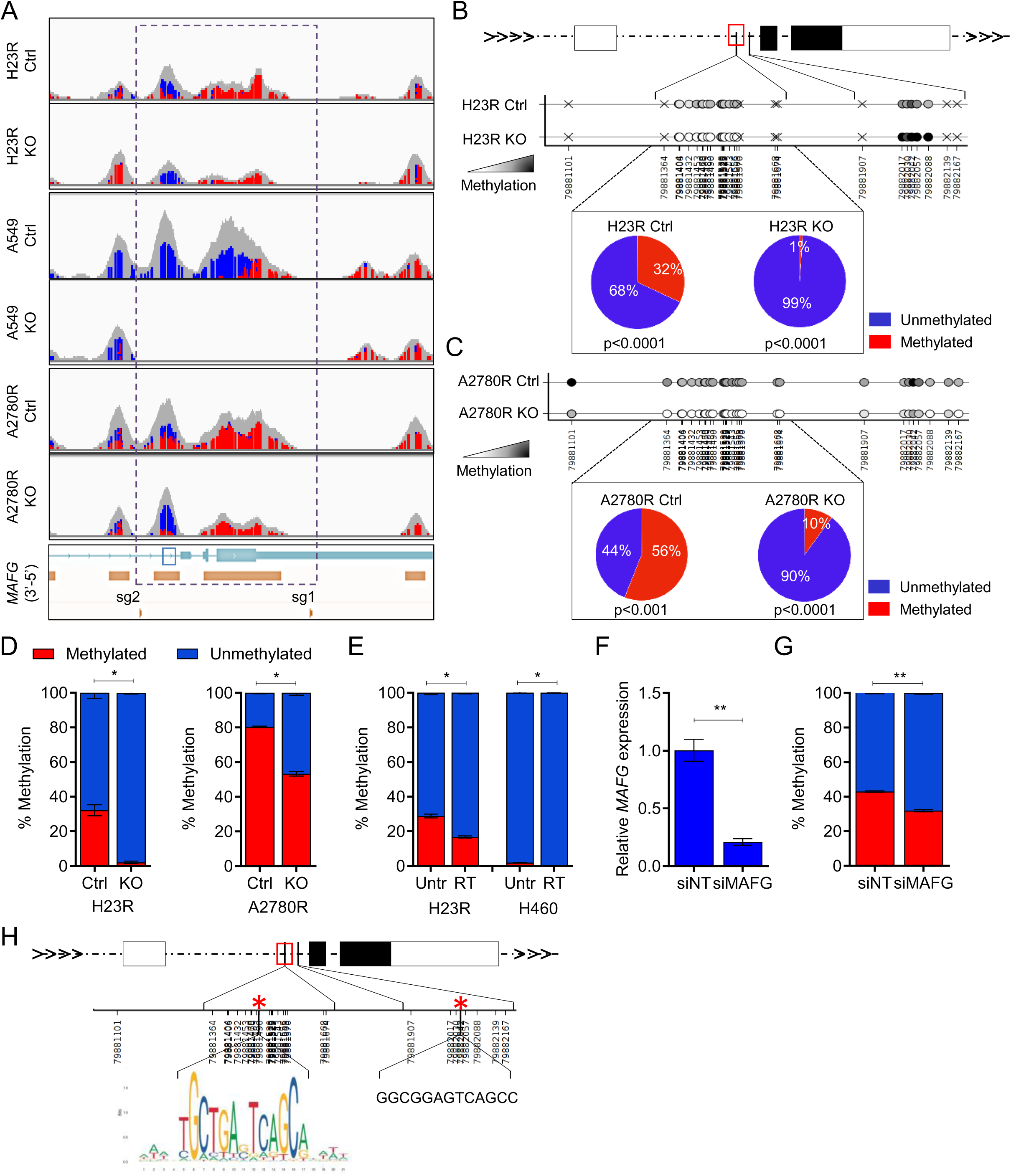
Epigenetic autoregulation of MAFG in lung cancer cell lines. **(A)** Representation of sequencing reads across the MAFG promoter region in control (Ctrl) and MAFG knockout (KO) clones from H23R, A2780R and A549 cell lines. Red indicates hypermethylated reads, and blue indicates hypomethylated reads. **(B,C)** Genomic position, methylation status, and relative proportions of methylated and unmethylated reads (pie charts) for 15 CpG sites shared between H23R (B) and A2780R (C) samples, located within a regulatory region of the MAFG gene (red box). **(D)** Validation of MAFG methylation changes in H23R and A2780R cell lines by quantitative methylation-specific PCR (qMSP). n = 2 technical replicates from one out of two biological replicates are represented. Statistical significance was determined using Welch’s two-tailed t-test using the Ctrl group as reference. Error bars represent mean + s.d. *p<0.05. **(E)** Percentage of MAFG methylation measured by qMSP in cisplatin-resistant cell lines (H23R and H460R) comparing untreated (Untr) with 5′-azacytidine treated cells (RT). n = 2 technical replicates from one out of two biological replicates are represented. Statistical significance was determined using Welch’s two-tailed t-test using the Ctrl group as reference. Error bars represent mean + s.d. *p<0.05. **(F)** Validation by qRT-PCR of the MAFG knockdown in H23R cells transfected with MAFG-specific siRNA (siMAFG) compared to non-targeting siRNA control (siNT). *GAPDH* was used as endogenous control. n = 3 technical replicates from one out of two biological replicates are represented. Statistical significance was determined using Welch’s two-tailed t-test using the Ctrl group as reference. Error bars represent mean + s.d. **p<0.01. **(G)** MAFG methylation levels measured by qMSP in H23R cells following siRNA-mediated MAFG silencing. n = 2 technical replicates from one out of two biological replicates are represented. Statistical significance was determined using Welch’s two-tailed t-test using the Ctrl group as reference. Error bars represent mean + s.d. **p<0.01. **(H)** Schematic representation of the location of the identified MARE sequences in the promoter regions of MAFG The red box highlights the intronic CpG island within MAFG showing differential methylation.

To determine whether the observed loss of methylation was attributable to MAFG-dependent epigenetic regulation rather than a secondary effect of CRISPR/Cas9-mediated genomic alteration, we next evaluated MAFG methylation in cisplatin-resistant lung cancer cell lines treated with the DNA demethylating agent 5′-azacytidine (RT). In both lung cancer H23R and H460R cells, demethylating treatment resulted in decreased DNA methylation at the MAFG promoter region, with a significant reduction observed in H23RT and H460RT cells compared to their untreated counterparts (**Figure 3E**), being the changes more noticeable in H23R. These findings indicate that the identified CpG region is susceptible to methylation-dependent regulation independently of CRISPR/Cas9 editing. To further confirm that MAFG itself contributes to this epigenetic regulation, we transiently silenced MAFG in parental H23R cells using MAFG-specific siRNAs. After validating efficient knockdown (**Figure 3F**), we detected a 12% reduction in MAFG methylation in siMAFG-treated cells compared with non-targeting controls (**Figure 3G**). Collectively, these results support the existence of a feedback mechanism in which MAFG influences the methylation state of its own regulatory region. To complement these results, we sought to confirm the potential DNA-binding regions of MAFG in its regulatory region. A 1,804-bp genomic fragment corresponding to the intron-1 CpG cluster of *MAFG* (hg19: chr17:79,880,864–79,882,667) was scanned using FIMO and the canonical MARE motif (TGCTGAGTCAGCA). The analysis revealed a high-affinity MARE site at positions 627–639 (p = 2.14 × 10^−7^), corresponding to the canonical sequence on the reverse strand and its palindromic counterpart on the forward strand. This site overlaps the CpG cluster that shows differential methylation in our models. A second, lower-affinity MARE-like site was detected at positions 1177–1189 (p = 1.45 × 10^−5^) (**Figure 3H**), altogether supporting the hypothesis that MAFG may bind and regulate its own regulatory region.

Following the identification of a methylation-sensitive CpG region associated with MAFG, we investigated its clinical relevance in NSCLC patients. As an initial step, we evaluated the suitability of digital PCR (dPCR) as an alternative to qMSP for methylation detection in patient samples. A positive correlation (R2: 0.69; p=0.0003) was observed between qMSP and dPCR measurements (**Supplementary Figure S4A**), supporting the use of dPCR for subsequent analyses. We then analyzed MAFG methylation in a cohort comprising 33 fresh-frozen (FT) and 37 FFPE (PT) samples from patients with early-stage NSCLC (**Table 1**). Methylation values from both sample types exhibited similar non-normal distributions (**Supplementary Figure S4B**) and did not differ significantly (**Supplementary Figure S4C**), justifying the combination of both sample types for survival analysis. Kaplan–Meier survival analyses revealed no significant association between MAFG methylation levels and progression-free survival (PFS) or overall survival (OS) in either lung adenocarcinoma (LUAD) or lung squamous cell carcinoma (LUSC) within this cohort (**Supplementary Figure S4D,E**). To further evaluate the biomarker potential of MAFG methylation, we extended our analysis to two larger TCGA NSCLC cohorts. We selected five CpGs within our region of interest (**Figure 4A** and **4B**) and patients were stratified based on average methylation levels. LUAD (n = 483; cut-off 0.4335) and LUSC (n = 364; cut-off 0.3657) cohorts were analyzed independently. In LUAD, low MAFG methylation was significantly associated with poorer prognosis for both PFS (p = 0.028) and OS (p = 0.0040) (**Figure 5C,D**). No significant associations were observed in the LUSC cohort (**Figure 5E,F**). Given previous reports supporting MAFG expression as a prognostic marker in lung cancer^6^, we next assessed whether MAFG protein levels also held prognostic value. Using two MAFG-specific protein aptamers (aptMAFG3F and aptMAFG6F)^6^ (**Figure 4G**), we analyzed MAFG protein expression in 127 FFPE samples from patients with early-stage NSCLC (**Table 1**). Survival analyses incorporating clinical-pathological variables showed no overall association between MAFG protein levels and OS when considering the combined cohort (**Figure 4H,I**). However, stratification by histological subtype revealed that higher MAFG levels detected using aptMAFG3F were significantly associated with worse OS in LUAD patients (p = 0.0225) (**Figure 4J**), with a similar trend for aptMAFG6F (p=0.0954) (**Figure 4K**). No significant associations were detected in LUSC patients using aptMAFG3F (p=0.1100) (**Figure 4L**), although higher MAFG levels detected by aptMAFG6F showed improved prognosis (p = 0.0367) (**Figure 4M**). No statistically significant associations were observed for PFS (**Supplementary Figure S4G-I**). Altogether, these findings demonstrate that MAFG is subject to methylation-dependent autoregulation and support a histology-specific clinical relevance of both MAFG methylation and protein expression, with prognostic significance restricted to lung adenocarcinoma. These results further suggest differential regulatory roles for MAFG in LUAD and LUSC.

**Figure 4.**
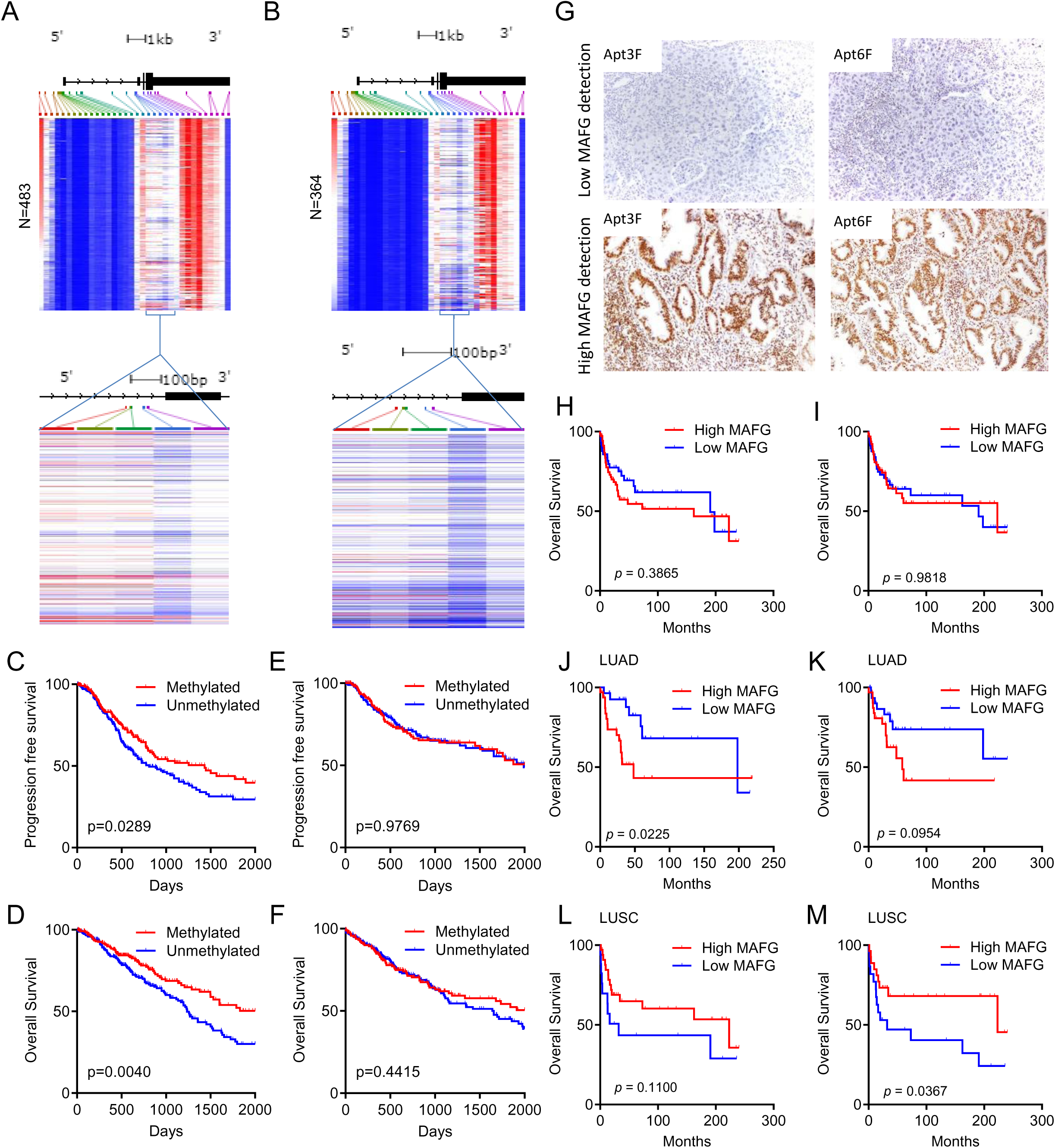
Clinical relevance of MAFG in NSCLC patients. **(A,B)** Methylation levels of MAFG-associated CpG islands in the TCGA LUAD (A) and LUSC (B) data available at https://xenabrowser.net/. **(C-F)** Kaplan–Meier analysis of progression-free survival (PFS) and overall survival (OS) in the TCGA LUAD (C,D) and LUSC (E,F) data based on methylation levels of CpG sites within the MAFG promoter from A and B using https://xenabrowser.net/. Cut-off Beta-value for LUAD is 0.4335 and for LUSC is 0.3657. MAFG Methylated in LUAD n = 240; MAFG Unmethylated in LUAD n = 243; MAFG Methylated in LUSC n = 167; MAFG Unmethylated in LUSC n = 198. Log-rank (Mantel-Cox) test was used to determine differences in survival between groups. **(G)** Representative aptahistochemistry images showing MAFG detection in FFPE tumor samples with low and high MAFG expression. **(H,I)** Kaplan–Meier analysis of overall survival (OS) in 127 early-stage NSCLC patients based on MAFG levels detected using AptMAFG3F (H) or AptMAFG6F (I). **(J-M)** Subgroup Kaplan–Meier analyses according to histological subtype, showing OS in LUAD (J,K) and LUSC (L,M) patients stratified by MAFG expression detected with AptMAFG3F (J,L) or AptMAFG6F (K,M). Median expression values were used as cut-off thresholds. Log-rank (Mantel-Cox) test was used to determine differences in survival between groups.

**Figure 5.**
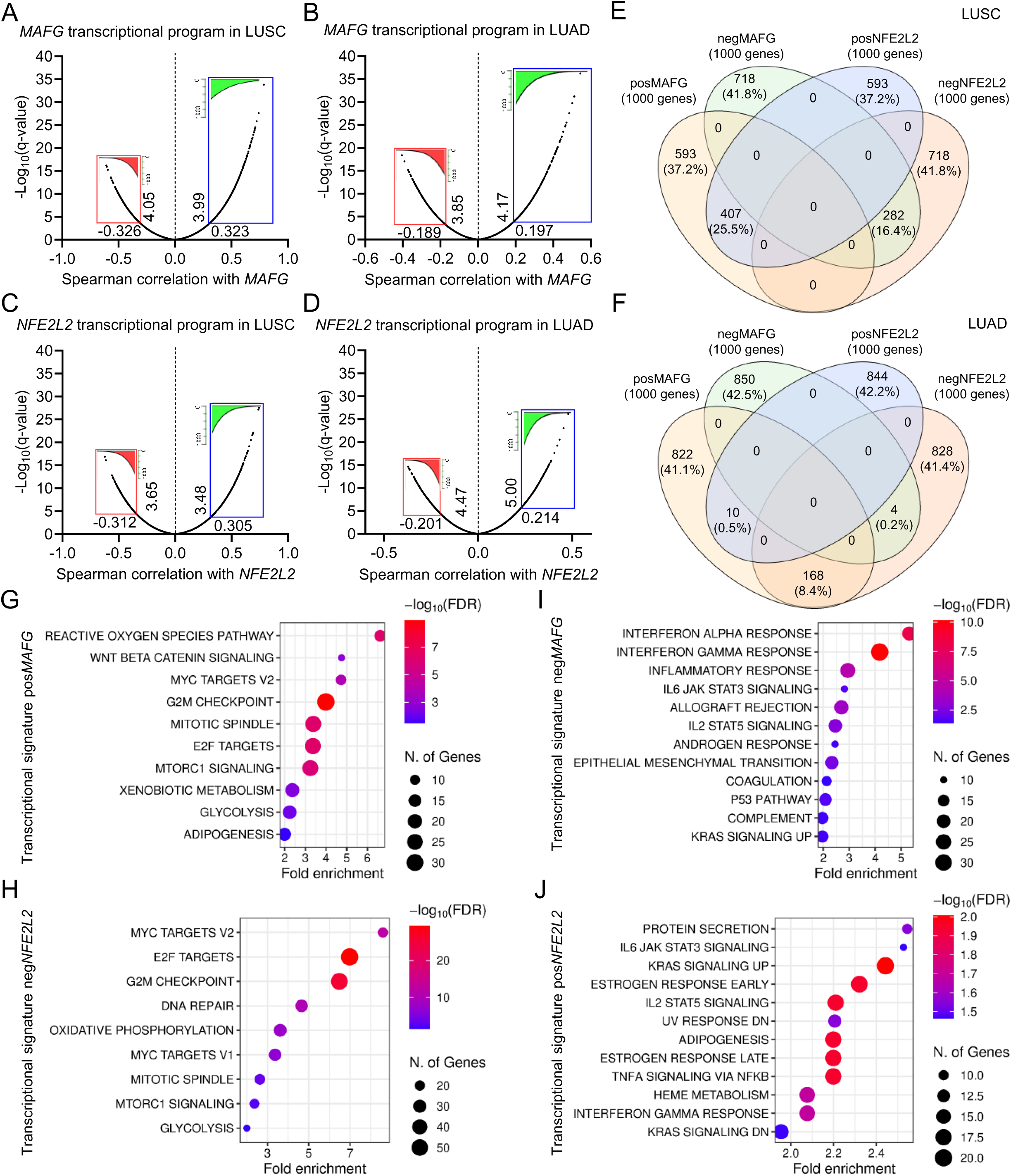
Distinct transcriptional programs associated with MAFG in LUAD and LUSC. **(A–D)** Transcriptional signatures associated with MAFG (A,B) and NFE2L2/NRF2 (C,D) in TCGA LUSC and LUAD cohorts. The top 1,000 genes showing the strongest positive (green) or negative (red) expression correlations are shown. **(E,F)** Venn diagrams overlapping the different transcriptional signatures selected in A-D for LUSC (E) and LUAD (F). **(G-J)** Molecular pathways enriched among genes associated with the positive transcriptional signatures of *MAFG* (G), the negative transcriptional signatures of *NFE2L2* (H), the negative transcriptional signatures of *MAFG* (I) and the positive transcriptional signatures of *NFE2L2* (J) in LUAD. Pathway enrichment analysis was performed using ShinyGO v0.85 (Ge SX, Jung D & Yao R, Bioinformatics 36:2628–2629, 2020) with the Hallmark gene sets from MSigDB and an FDR cutoff of 0.05.

**Table 1:**
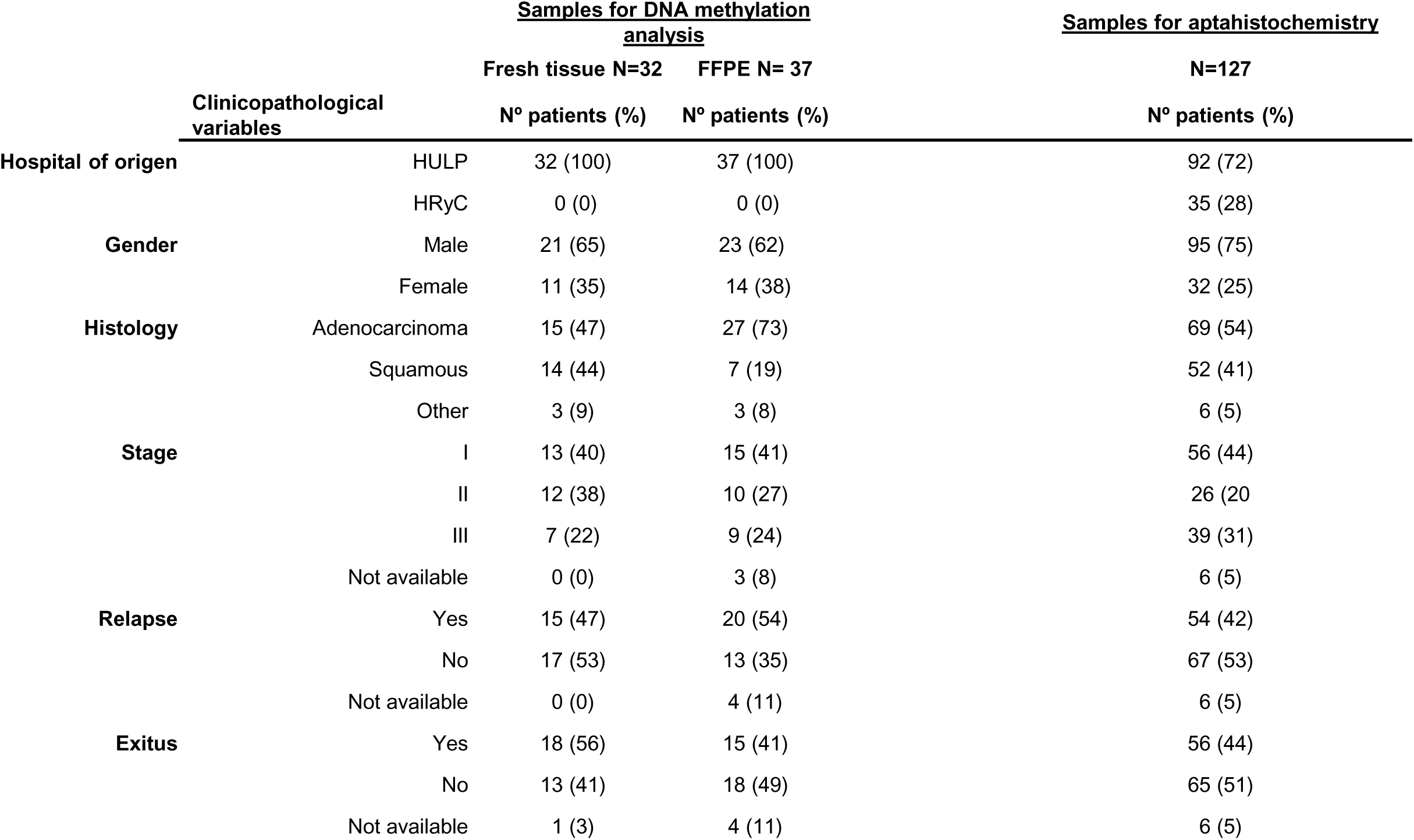
Demographic data on patients with early-stage NSCLC at HULP used in this study.

|  |  | <u>Samples for DNA methylation analysis</u> |  | <u>Samples for aptahistochemistry</u> |
| --- | --- | --- | --- | --- |
|  |  | Fresh tissue N=32 | FFPE N= 37 | N=127 |
|  | Clinicopathological variables | N° patients (%) | N° patients (%) | N° patients (%) |
| Hospital of origen | HULP | 32 (100) | 37 (100) | 92 (72) |
|  | HRyC | 0 (0) | 0 (0) | 35 (28) |
| Gender | Male | 21 (65) | 23 (62) | 95 (75) |
|  | Female | 11 (35) | 14 (38) | 32 (25) |
| Histology | Adenocarcinoma | 15 (47) | 27 (73) | 69 (54) |
|  | Squamous | 14 (44) | 7 (19) | 52 (41) |
|  | Other | 3 (9) | 3 (8) | 6 (5) |
| Stage | I | 13 (40) | 15 (41) | 56 (44) |
|  | II | 12 (38) | 10 (27) | 26 (20) |
|  | III | 7 (22) | 9 (24) | 39 (31) |
|  | Not available | 0 (0) | 3 (8) | 6 (5) |
| Relapse | Yes | 15 (47) | 20 (54) | 54 (42) |
|  | No | 17 (53) | 13 (35) | 67 (53) |
|  | Not available | 0 (0) | 4 (11) | 6 (5) |
| Exitus | Yes | 18 (56) | 15 (41) | 56 (44) |
|  | No | 13 (41) | 18 (49) | 65 (51) |
|  | Not available | 1 (3) | 4 (11) | 6 (5) |

### MAFG is associated with distinct transcriptional programs in LUAD and LUSC

The differential prognostic impact of MAFG observed between lung adenocarcinoma and lung squamous cell carcinoma suggests that MAFG may exert subtype-specific molecular functions. To investigate this possibility, we compared the transcriptional programs associated with MAFG expression in these two NSCLC subtypes. MAFG is classically described as a transcriptional partner of NRF2 (encoded by NFE2L2), a master regulator of oxidative stress responses and a well-established oncogenic driver in lung cancer^15^. To contextualize MAFG-associated transcriptional changes, we therefore included genes correlated with *NFE2L2* expression as a reference for canonical NRF2-driven pathways. Using TCGA datasets, we identified the top 1,000 genes showing positive or negative correlation with *MAFG* or *NFE2L2* expression in LUAD (n = 587) and LUSC (n = 501) cohorts (**Figure 5A**-**5D**).

Comparison of these gene sets revealed marked differences between the two histological subtypes. In LUSC, *MAFG* and *NFE2L2* showed substantial overlap, sharing positively correlated gene expression in 25.5% of genes and negatively correlated expression in 16.4% (**Figure 5E**). In contrast, overlap between *MAFG*- and *NFE2L2*-associated genes in LUAD was minimal, with only 0.5% of positively correlated genes and 0.2% of negatively correlated genes shared (**Figure 5F**). Moreover, analysis of *MAFG* and *NFE2L2* transcriptional signatures revealed 168 genes (8.4%) that were positively correlated with MAFG but negatively correlated with *NFE2L2* in LUAD, a pattern that was not observed in LUSC (**Figure 5E,F**). Pathway enrichment analysis further highlighted these differences. In LUSC, both *MAFG* and *NFE2L2* positively correlated with pathways related to cytotoxic and oxidative stress responses, including reactive oxygen species and xenobiotic metabolism (**Supplementary Figure S5A**), with pathways such as epithelial-mesenchymal transition, inflammatory response and hypoxia negatively associated to both *MAFG* and *NFE2L2* (**Supplementary Figure S5B**). In contrast, in LUAD, cytotoxic response pathways were positively associated with *MAFG* but showed limited or no association with *NFE2L2* (**Figure 5G**). Notably, cell cycle–related pathways (including MYC targets, G2/M checkpoint, mitotic spindle, E2F targets, and mTORC1 signaling) were also positively correlated with *MAFG* (**Figure 5G**) and negatively correlated with *NFE2L2* expression in LUAD (**Figure 5H**), whereas inflammatory and immune-related pathways (inflammatory response, IL2/STAT5 signaling, and KRAS signaling) displayed the opposite pattern (**Figure 5I-J**). Together, these results indicate that while *MAFG* and *NRF2* are transcriptionally aligned in LUSC, MAFG is associated with a distinct, and in some cases opposing, transcriptional program in LUAD. This suggests that, in lung adenocarcinoma, MAFG may act outside the canonical NRF2-dependent antioxidant network and instead engage pathways related to proliferation and oncogenic signaling.

To further explore the broader functional implications of MAFG-dependent epigenetic regulation in LUAD, we performed Gene Ontology and pathway enrichment analyses using three independent gene sets: (i) the 56 genes exhibiting methylation changes following MAFG KO in our study (GO1; **Supplementary Figure S2A**), (ii) the 83 genes previously validated after MAFG knockdown in colorectal cancer and melanoma by Fang et al.^12,13^ (GO2, **Figure 6A**), and (iii) the top 1,000 genes negatively correlated with MAFG expression in the TCGA LUAD cohort (GO3, **Figure 5I**). Several pathways, including xenobiotic metabolism, inflammatory response, and IL2/STAT5 signaling, were shared by at least two of the three gene sets. Notably, the only pathway common to all three was “KRAS signaling UP” (**Figure 6B**), which included 54.5% of well-established tumor suppressor genes, 22.7% oncogenes and 22.7% of context-dependent genes (**Supplementary Table 2**). Given the clinical relevance of KRAS alterations in lung adenocarcinoma, we next evaluated the prognostic impact of MAFG expression and methylation in LUAD patients harboring KRAS gain-of-function mutations within the TCGA cohort. Among patients with confirmed KRAS mutations (n = 23), high MAFG expression (p = 0.0300, **Figure 6C**) and low MAFG methylation (p = 0.0055, **Figure 6D**) were significantly associated with shorter progression-free survival. To increase statistical power, we performed an expanded analysis selecting the top 25% of LUAD patients with highest KRAS expression (n = 129). Consistently, higher MAFG expression (p = 0.0406, **Figure 6E**) and lower methylation levels (p = 0.0043, **Figure 6F**) were associated with poorer progression-free survival in this group. Altogether, these results support a model in which MAFG contributes to a LUAD-specific transcriptional program that is partially decoupled from canonical NRF2 signaling and is closely linked to KRAS-associated oncogenic pathways. This provides a mechanistic framework to explain the subtype-specific prognostic value of MAFG observed in lung adenocarcinoma but not in lung squamous cell carcinoma.

**Figure 6.**
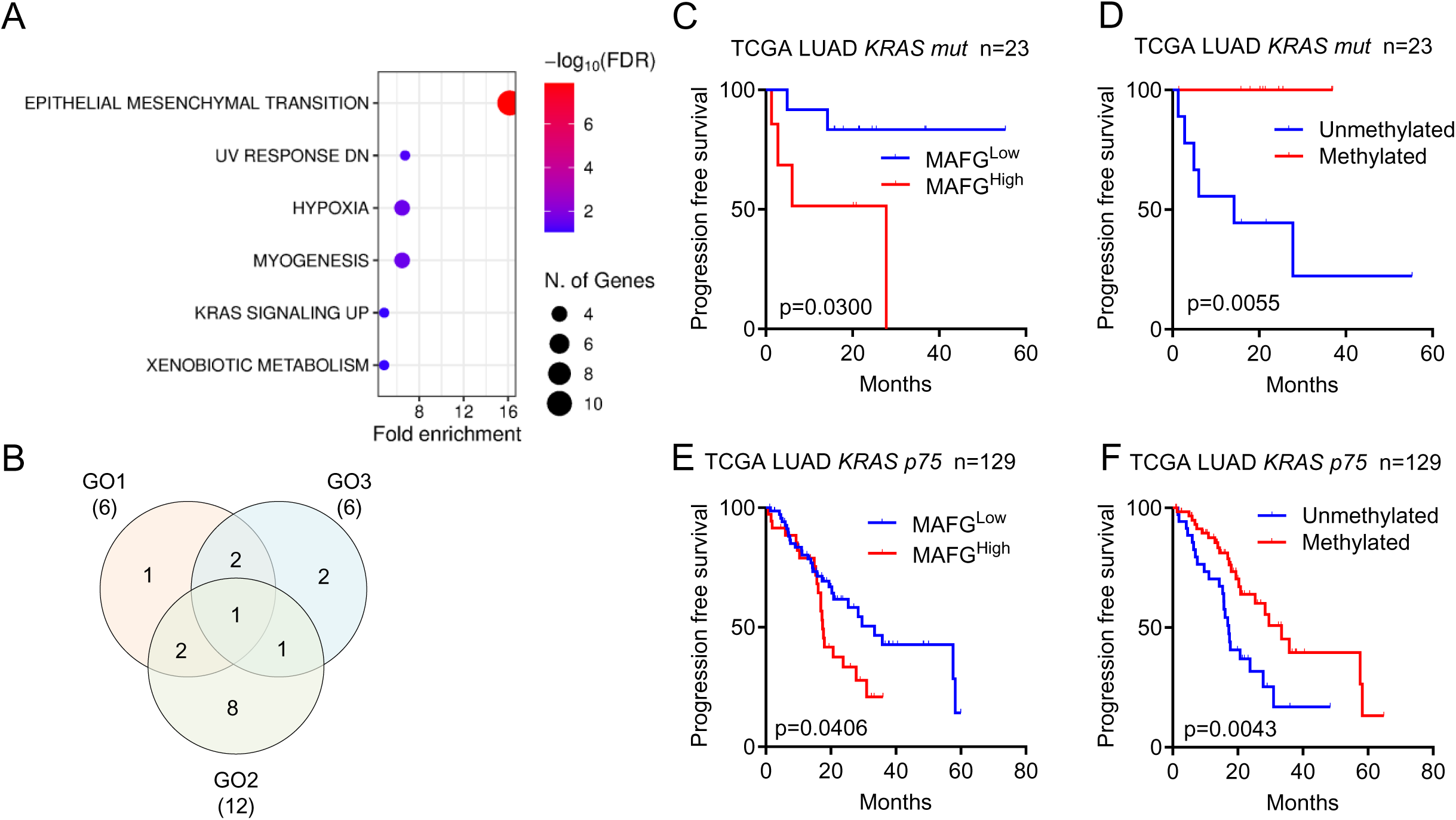
Novel clinical implications of MAFG in KRAS-mutant lung adenocarcinoma (LUAD) patients. **(A)** Gene Ontology analysis with genes whose expression changes following MAFG knockdown in colorectal cancer and melanoma (GO3), obtained from Fang et al. 2014 and 2016. Pathway enrichment analysis was performed using ShinyGO v0.85 (Ge SX, Jung D & Yao R, Bioinformatics 36:2628–2629, 2020) with the Hallmark gene sets from MSigDB and an FDR cutoff of 0.1. **(B)** Venn diagram showing overlap among molecular pathways identified by GO analysis using GO1 (Pathway enrichment analysis of the 56 differentially methylated genes identified by SeqCap after MAFG deletion from Figure S2A), GO2 (Pathway enrichment analysis of the 1000 negatively correlated genes from Figure 5I) and GO3 from Figure 6A. **(C,D)** Kaplan–Meier analyses of progression-free survival in the TCGA cohort of 23 LUAD patients harboring gain-of-function KRAS mutations, stratified according to MAFG expression (C) or MAFG methylation (D). **(E,F)** Kaplan–Meier analyses of progression-free survival in the top 25% of TCGA patients with high KRAS expression (p75, n = 129), stratified according to MAFG expression (E) or MAFG methylation (F). Log-rank (Mantel-Cox) test was used to determine differences in survival between groups.

## DISCUSSION

The search for new biomarkers to predict and monitor cancer patients is crucial for clinical decision-making, especially in lung cancer, where the 5-year survival rate for NSCLC patients is 26.4%^16^. Previous studies suggest that MAFG is a potential biomarker for chemotherapy resistance in NSCLC patients. However, whether its mechanism of action is through direct transcription or epigenetic silencing is still unknown. Here we show that MAFG is a pro-oncogenic factor that can be self-regulated by DNA methylation and which different regulation layer can be used as prognostic biomarker in lung adenocarcinoma patients.

To characterize the methylation changes driven by MAFG, we established a MAFG knock-out in vitro model using CRISPR/Cas9 technology in three epithelial cancer cell lines. Successful MAFG deletion was confirmed for all three cell lines, being homozygous in lung cancer cell lines H23R and A549, and heterozygous in ovarian cancer cell line A2780R. These results, together with existing literature, suggest uneven sMAFs distribution among tissues and compensatory mechanisms in lung cancer cells that does not occur in the ovarian cancer cell line of this study. Despite MAFG KO mice showing mild thrombocytopenia and motor ataxia that lessens with age^17–19^, these studies support that MAFG is dispensable for development and that its global loss is well tolerated.

We then characterized the epigenetic regulation driven by MAFG using SeqCap technology, which analyzes 10 times more CpGs than the HM450k methylation data array^20,21^. We identified 56 candidate genes with significant methylation changes, prioritizing those present in 4 out of 5 analyses: *ADRB1*, *LIF*, miR-4456, and *MAFG* itself. Among the 56 candidate genes we identified *DUSP1*, which was included in the final candidate selection due to its significant involvement in cancer^22,23^. Transcriptional validation suggested that only *DUSP1* and *LIF* methylation changes after MAFG KO altered their transcription. Additionally, in a small cohort of 10 NSCLC patients (5 LUAD and 5 LUSC), we found a negative correlation of *LIF* and *MAFG* in both LUAD and LUSC patients and for *DUSP1* and *MAFG* in LUAD patients. Despite the relevance of *DUSP1* methylation in cancer^24–27^, our SeqCap analysis showed only 4 hypomethylated CpGs in the MAFG KO group out of 25 positions identified in its promoter region, leading us to discard *DUSP1* for additional validations.

Using SeqCap and qMSP, we identified and confirmed the hypomethylation of several CpGs in the *LIF* promoter region in A549 cells lacking MAFG. Some studies indicate that LIF/LIFR is under epigenetic regulation in cancer with changes *LIF* methylation described in meningioma^28^ and breast cancer^29^. Thus, we investigated translational implications of *LIF* methylation in NSCLC. By analyzing the association between methylation and disease progression in the TCGA we did not observe a significant prognostic role for the methylation in the positions analyzed. The TCGA methylation data for *LIF* did not cover all 5′ positions of the promoter region identified in our study, which may explain discrepancies in the epigenetic regulation we observed and its potential implications for patient outcomes. Therefore, examining additional *LIF* promoter CpGs is necessary to better define its relevance in NSCLC.

The most surprising finding was the identification of a potential epigenetic self-regulation of *MAFG*. Specifically, we found a region of 15 CpG positions in a CpG island near the end of intron 1 of *MAFG*, which loses methylation in MAFG KO groups of H23R and A2780R. Validation by qMSP in both cell lines, together with the observation that 5-Aza treatment reduces methylation levels of the analyzed region in H23R, and the effect of *MAFG* silencing over its methylation levels, confirmed that *MAFG* is under epigenetic regulation. Consistent with this, previous studies have described a MARE sequence in the promoter of the antisense gene *MAFG-AS1*, enabling its transcriptional regulation by *MAFG* in bladder cancer^7^. Another recent report linked *MAFG-AS1* methylation to ovarian cancer dissemination^30^. Because both genes share the same promoter region, similar DNA-level regulatory mechanisms are likely involved. Indeed, our *in silico* analysis of the *MAFG* promoter identified two MARE motifs within its regulatory region, which, together with our experimental data, strongly supports the existence of *MAFG* self-regulation. Further studies using intermediate levels of *MAFG* silencing will be required to define the dynamics of this mechanism and clarify the role of *MAFG* as a modulator of DNA methylation.

Regarding its clinical use, few studies report a relationship between *MAFG* overactivation and tumor aggressiveness in hepatocellular carcinoma^10^ or lung cancer^6^. Additional reports founds an association between MAFG high levels and worse prognosis in NSCLC patients^6^, melanoma, breast cancer, LUSC, or hepatocellular carcinoma^8^. However, there is no study linking *MAFG* promoter methylation or protein levels to clinical outcomes in any cancer subtype. Thus, we analyzed the methylation levels of *MAFG* in a pilot cohort of 70 NSCLC patients using digital PCR, and their association with clinical-pathological characteristics. Due to the reduced sample size and assay specificity, we combined fresh and FFPE samples but we did not observe a significant association between methylation levels and the disease progression. To further investigate this, we studied *MAFG* methylation levels in the lung adenocarcinoma (LUAD) and lung squamous cell carcinoma (LUSC) cohorts of the TCGA, analyzed by HM450k. Higher *MAFG* methylation levels were associated with better prognosis in LUAD but not in LUSC patients. These results suggest a potential role for *MAFG* as a biomarker in LUAD. Since the methodology used for the analysis was different -TCGA data are based on sequencing reads from the HM450k methylation panel, our data are based on methylation probes measured by digital PCR-, we cannot exclude that different sensitivities of each technique can account for the different results.

We also evaluated MAFG protein levels as a prognostic marker in NSCLC. We studied the relationship between MAFG protein expression and survival and clinical-pathological data in 127 paraffin-embedded tumor tissue samples from patients with early-stage NSCLC. Using specific MAFG aptamers identified in a previous work (ApMAFG3F and ApMAFG6F)^6^, we found that higher detection of MAFG by ApMAFG3F, and to a lesser extent ApMAFG6F, predicts worse clinical prognosis in LUAD patients. Interestingly, we observed an opposite trend in LUSC patients, suggesting different implications of MAFG between LUAD and LUSC, as seen in the methylation analysis.

These observations lead us to hypothesize that MAFG may have different molecular functions between LUAD and LUSC subtypes in NSCLC. Given the role of MAFG as a transcription factor activating the antioxidant response with NRF2, we analyzed the transcriptional signature associated with *MAFG* and *NFE2L2* in these two TCGA cohorts. We found that LUSC patients shared a higher percentage of genes in each transcriptional signature, whereas the shared transcriptional signature between *MAFG* and *NFE2L2* in LUAD patients is much lower, suggesting different transcriptional regulation related to the cytotoxic response between the subtypes. These differences might also be explained by the incidence of mutations in *KEAP1* and *NFE2L2* reported for LUAD and LUSC. While mutations in *KEAP1* and *NFE2L2* are 12.1% and 14.9% for LUSC patients, respectively, in LUAD 17.1% of patients have *KEAP1* mutations and 2.2% have *NFE2L2* mutations. These mutations are mutually exclusive, favoring constitutive NRF2 activation^31–33^. Interestingly, only in LUAD transcriptional signatures we found genes positively associated with *MAFG* and negaively with *NFE2L2*. Additionally, gene ontology analyses showed that there are molecular pathways that have an opposite correlation with *MAFG* and *NFE2L2* in LUAD. Previous reports described a subpopulation of astrocytes with a MAFG^+^/NFE2L2^−^ transcriptional phenotype, where MAFG silences NRF2 target genes to promote the activation of pro-inflammatory genes^14^. Although the role of MAFG in regulating pro-inflammatory processes in LUAD is unknown, it might negatively regulate NRF2 transcriptional signature, not only as a transcriptional repressor but also as a methylating agent. Further studies will be required to define the precise molecular relationship between MAFG and the epigenome in LUAD patients.

Additionally, the ontological analysis of the 56 candidate with significant hypomethylation in the *MAFG* KO group showed associations with the cytotoxic response, KRAS signaling, and inflammatory and cytokine. By combining this analysis with two additional transcriptional signatures, the 87 genes epigenetically silence by *MAFG* in CRC and melanoma^12,13^, and the transcriptional signature contrary to *MAFG* in the LUAD cohort of TCGA, we found KRAS signaling as a common hallmark in all the three approaches. Most of the genes in this pathway that negatively correlate with MAFG in our analysis correspond to well-established tumor suppressors, including key negative regulators of the RAS/MAPK pathway (DUSP6, SPRY2, MAP3K1)^34–36^, mediators of apoptosis or senescence (IGFBP3, GNG11, FUCA1)^37–39^, and genes involved in epithelial integrity (PIGR, TFP)^40,41^. This pattern is consistent with MAFG-high tumors adopting a KRAS-rewired transcriptional state in which classical KRAS-UP genes are attenuated, potentially through MAFG-dependent epigenetic remodeling. Importantly, KRAS is highly overactivated in ∼30% adenocarcinoma subtypes like pancreatic cancer, CRC or NSCLC^42–44^, but not in squamous subtypes, like LUSC, where only a 4% of *KRAS* mutations have been reported^44^. Analyzing the status of *MAFG* in a small subgroup of patients from the global TCGA adenocarcinoma cohort with gain-of-function KRAS mutation, we observed that both higher methylation and low expression of *MAFG* could be markers of good prognosis for disease progression. This significance is maintained even in patients with high *KRAS* expression. These results indicate that *MAFG* methylation is a promising biomarker of good prognosis in NSCLC adenocarcinoma, especially in those with *KRAS* mutation.

Altogether, these results demonstrate that *MAFG* plays a central role in shaping the epigenetic landscape of NSCLC and support its relevance as a molecular target for future preclinical exploration. Moreover, the translational evidence presented here, together with previous studies, indicates that detection of *MAFG* at the DNA methylation, mRNA expression, and protein levels constitutes a promising biomarker for NSCLC patients, particularly within the LUAD subtype.

## METHODS

### Cell lines

Three immortalized human cell lines (ATCC) were used in this study: H23 and A549 non-small cell lung cancer (NSCLC) cell lines, and A2780 ovarian cancer cells. Cisplatin-resistant sublines (R) of the H23 and A2780 cell lines were previously generated in our laboratory^45^. All cell lines were maintained in RPMI-1640 medium (Roswell Park Memorial Institute) supplemented with 10% FBS (Thermofisher, USA; CAT. #15377636), L-glutamine (Gibco, USA; CAT #25030- 024), gentamicin (Gibco, USA; CAT #REF 15750-078) and amphotericin B (Corning, USA; REF 30-003-CF), and cultured at 37°C in a humidified atmosphere containing 5% CO_2_. HEK293T cells were obtained from ATCC and cultured in DMEM containing 10% FBS and 1% penicillin/streptomycin at 37°C in a humidified atmosphere containing 5% CO_2_. All cell lines were routinely tested for mycoplasma using MycoAlert Plus (Lonza, Suiza; CAT #LT07-710), and human melanoma cell lines were STR authenticated by the Genomics Core at Instituto de Investigaciones Biomédicas Alberto Sols.

### Clinical samples

A total of 196 NSCLC tumor tissue samples were included in this study. All patients were diagnosed with early-stage NSCLC (stages I–III) at Hospital Universitario La Paz and underwent complete surgical tumor resection. All cases presented with a positive positron emission tomography (PET) scan showing localized lung tumors and were histologically confirmed by the Pathology Department of Hospital Universitario La Paz. Written informed consent was obtained from all patients, and samples were requested through the Biobank of Hospital Universitario La Paz in accordance with institutional and ethical guidelines. Samples were distributed as follows: 32 fresh-frozen NSCLC tumor specimens were stored at −80 °C from surgery until laboratory processing, and 37 formalin-fixed paraffin-embedded (FFPE) tumor samples were stored at the HULP Biobank. Both sample sets were used for DNA extraction and subsequent methylation analysis by quantitative methylation-specific PCR (qMSP). Additionally, 10 fresh tumor samples were used for mRNA expression analyses. Furthermore, 127 FFPE tumor samples were sectioned using a microtome, mounted onto glass slides, and used for aptamer-based immunohistochemical analyses targeting MAFG. Demographic and clinicopathological characteristics of the patient cohorts are summarized in Table 1.

### CRISPR/Cas9-mediated gene editing of MAFG

Deletion of the MAFG gene was achieved using CRISPR/Cas9 technology through two independent delivery approaches. To generate a complete removal of the MAFG coding sequence, two single-guide RNAs (sgRNAs) targeting distinct regions of the MAFG locus were designed using multiple bioinformatic tools (https://www.zlab.bio/resources and https://www.benchling.com/). sgRNAs with the highest predicted specificity and lowest off-target probability were selected: sgMAFG-1: 3′-CCAGAGCTAGATGGACGGCCCAC-5′ (PAM: CCA); sgMAFG-2: 5′-TGTGCGCTTAACGTGAGGG-3′ (PAM: GGG). Recombinant Cas9 protein and sgRNAs were obtained from IDT Technologies (USA).

#### Electroporation

Electroporation was performed in collaboration with the Molecular Cytogenetics Unit at the Spanish National Cancer Research Centre (CNIO). Cas9 protein complexes with sgRNAs were delivered into A549 and A2780R cells using the Neon Transfection System (Thermo Fisher Scientific, USA; CAT #MPK5000). Control (Ctrl) cells were electroporated with Cas9 protein alone. Cells were harvested at approximately 75% confluence using trypsin, washed twice with PBS and resuspended in 10 µL of Buffer R per experimental condition. Approximately 200,000 cells per reaction were electroporated using two pulses of 1200 V for 30 ms. Cells were then seeded into 12-well plates containing pre-warmed RPMI medium. Subsequently, cells were plated into 96-well plates using serial dilution to enable single-cell clonal expansion and selection of successfully edited MAFG-KO clones.

Because clonal expansion of the H23R cell line was not successful using electroporation, lentiviral transduction was selected as an alternative approach due to its higher efficiency and lower associated cytotoxicity.

#### Lentiviral transduction

Lentiviral gene editing was performed in collaboration with the Translational Research Group in Head and Neck Cancer and Maxillofacial Surgery at IdIPaz, following a previously described protocol^46^. For Cas9 expression, HEK293T cells were transfected with the lentiCas9-Blast plasmid (Addgene plasmid CAT #52963) using polyethyleneimine (Polysciences, USA; CAT #24765). For sgRNA delivery, cells were transfected with a pLKO vector encoding MAFG-targeting sgRNAs, together with the lentiviral packaging plasmids pMD2-VSVg and pPAX2 (Addgene plasmids #12259 and #12260, respectively), using PEI. sgRNAs targeting dTomato were used as editing controls. Viral supernatants were collected at 48 h and 72 h post-transfection, filtered through 45 µm PVDF filters (Avantor, USA; CAT #MILFSLHV033NB), and used for infection. For MAFG editing, 300,000 H23R cells per well were seeded and incubated with a 1:2 dilution of viral supernatant in the presence of 8 µg/mL polybrene. Cells were first transduced with Cas9-expressing lentivirus and selected with blasticidin (5 µg/mL; Sigma Aldrich, USA; CAT. #15205) for 3 days. Subsequently, cells were transduced with sgRNA-expressing lentivirus and selected with puromycin (1 µg/mL; Sigma-Aldrich, USA; CAT #54011) for an additional 3 days. In both genome-editing approaches, MAFG deletion in the selected clones was confirmed at the DNA level by PCR, at the transcript level by quantitative PCR (qPCR), and at the protein level by Western blot (WB).

### Genomic DNA extraction and semi-quantitative PCR analysis

Genomic DNA was extracted from cell line samples and fresh tumor tissue samples by adding lysis buffer (87.5% DNA buffer B, 10% SDS, and 2.5% Proteinase K), followed by incubation at 37 °C for 24 h and subsequent phenol–chloroform digestion, as previously described^5^. DNA from formalin-fixed paraffin-embedded (FFPE) patient tissue samples was deparaffinized using xylene prior to extraction. DNA concentration and purity were assessed using a NanoDrop ND-1000 spectrophotometer (ThermoFisher, USA). Extracted DNA samples were stored at −20 °C until further use. For the selection of candidate cell clones following CRISPR/Cas9 gene editing, genomic DNA was precipitated using 5 M NaCl (0.015%) and absolute ethanol. Edited regions of the MAFG gene were amplified using two specific primer pairs (**Supplementary Table 3**) in combination with 10X Standard

Reaction Buffer (Biotools, Spain; CAT #10002-4100), using 500 ng of genomic DNA per reaction under the following PCR conditions: (a) 1 cycle of 5 min at 94 °C; (b) 45 cycles of 30 s at 94 °C, 30 s at 53 °C (primer annealing temperature), and 1 min 50 s at 72 °C; (c) final extension of 5 min at 72 °C. PCR products were resolved by electrophoresis on 1.5% agarose gels stained with SYBR Safe DNA Gel Stain (Invitrogen, USA; CAT #S33102). A 100-bp DNA ladder (New England Biolabs; USA; CAT #N3231S) was used to estimate amplicon size. Amplified fragments were excised and purified using the MiniElute Gel Extraction Kit (Qiagen, Germany; CAT #28604) and subsequently validated by Sanger sequencing to confirm MAFG gene editing.

### Methylome sequencing by Methyl-Seq and bioinformatic analysis

A total of 1,000 ng of genomic DNA was extracted from each cell line using conventional digestion methods followed by phenol-chloroform purification, as previously described^45^. Two independent biological replicates were generated for each experimental group (H23R Ctrl, H23R KO-MAFG, A2780R Ctrl, A2780R KO-MAFG, A549 Ctrl, and A549 KO-MAFG). DNA concentration and quality were assessed using a Qubit fluorometer (Thermo Fisher Scientific, USA; CAT #Q33226). Samples were submitted to GenomeScan (Leiden, The Netherlands) for bisulfite sequencing. Briefly, genomic DNA was fragmented to an average size of 300 bp and subjected to bisulfite conversion using the EZ-96 DNA Methylation-Lightning™ MagPrep Kit (Zymo Research, USA; CAT #D5031). Sequencing libraries were constructed using the NEBNext Ultra II DNA kit and double-stranded DNA adapters (IDT, USA), followed by target enrichment using the SeqCap Epi Hybridization (SeqCap) protocol (Roche)^20^. Sequencing was performed using the Illumina NovaSeq 6000 platform (Illumina). Sequencing data were analyzed in-house by our bioinformatics specialist Carlos Rodríguez Antolín. For each sample, a minimum of 80 million clusters and approximately 20 Mb of sequencing output were obtained. On-target coverage exceeded 30× depth for at least 80% of targeted positions, representing the recommended minimum quality threshold. Individual CpG sites covered at less than 10× depth were excluded from downstream analyses. The hg19 human genome assembly was used as the reference for data annotation. CpG methylation data were analyzed both individually for each cell line (H23R, A2780R, and A549) and in combined analyses, including a lung cancer-specific analysis (Lung: H23R + A549) and a global epithelial analysis (Epithelial: H23R + A2780R + A549). Data overdispersion was taken into account^47^, and statistical significance was assessed using Chi-square tests^48^. False discovery rates (FDR) were corrected using the Benjamini–Hochberg method^49^.For each analytical comparison, only CpG positions displaying significant methylation changes between cisplatin-resistant control (R Ctrl) and MAFG-knockout (R KO-MAFG) groups were considered.

### Bisulfite modification and quantitative methylation-specific PCR

Genomic DNA was subjected to sodium bisulfite treatment to preserve DNA methylation marks for subsequent quantification by quantitative methylation-specific PCR (qMSP). For cell line–derived samples, 1,000 ng of genomic DNA were denatured at 37 °C for 10 min and manually modified by treatment with hydroquinone (0.0011 g/mL) and sodium bisulfite (0.38 g/mL) at 50 °C for 17 h. Modified DNA was subsequently purified using the Wizard DNA Clean-Up Kit (Promega, USA; CAT #A7280). After purification, DNA samples were incubated with NaOH (0.3 mol/L) for 5 min at room temperature and precipitated using glycogen, ammonium acetate, and ethanol for 24 h at −20 °C. Additionally, genomic DNA from cisplatin-resistant (R) cell lines treated with the demethylating agent 5-azacytidine (5′-aza) (R-RT)^50^ was manually bisulfite-converted (1,000 ng) and MAFG methylation levels were quantified by qMSP. Genomic DNA obtained from NSCLC patient tumor samples was bisulfite-converted using the EZ DNA Methylation-Lightning Kit (Zymo Research, USA; CAT #D5031), following the manufacturer’s protocol. For quantitative methylation-specific PCR (qMSP), we designed primers and probes to evaluate DNA methylation of MAFG and LIF CpGs (**Supplementary Table 3**). qMSP reactions were performed in duplicate using the QuantiTect Multiplex PCR Kit (Qiagen, Germany; CAT #204443) and the StepOnePlus™ Real-Time PCR System (Roche, Switzerland) under the following conditions: (a) 1 cycle of 15 min at 95 °C; (b) 40 cycles of 1 min at 95 °C followed by 1 min at 60 °C. The percentage of DNA methylation for each sample was calculated using the previously described formula 100/ (1+(2^ (Ct_M – Ct_U)))^51^.

### DNA methylation analysis by digital PCR (dPCR)

MAFG DNA methylation levels in patient samples were quantified using the QuantStudio Absolute Q Digital PCR System (Applied Biosystems, Thermo Fisher Scientific, USA). A total of 69 tumor tissue samples from early-stage NSCLC patients were analyzed, including 32 fresh-frozen samples and 37 FFPE samples. The same primer pairs and methylated/unmethylated TaqMan probes used for qMSP (**Supplementary Table 4**) were used. Reactions were performed using the Absolute Q RT 1-Step Master Mix (Thermo Fisher Scientific, USA; CAT #A55165). For fresh-frozen tissue samples, a final bisulfite-modified DNA volume of 3 µL in a total reaction volume of 9 µL was used, with the following cycling conditions: 10 s at 96 °C, followed by 40 amplification cycles of 5 s at 96 °C and 30 s at 60 °C. For FFPE samples, 6 µL of bisulfite-modified DNA was used in a final volume of 9 µL, with cycling conditions of 10 min at 96 °C, followed by 40 amplification cycles of 10 s at 96 °C and 30 s at 60 °C. The percentage of DNA methylation for each sample was calculated using the formula: (copies of FAM / (copies of FAM + copies of VIC))*100

### RNA extraction and qRT-PCR analysis

Total RNA was extracted from both cell lines and fresh patient tissue samples using the guanidinium thiocyanate method with TRIzol™ Reagent (Thermofisher, USA; REF; am9738l) followed by chloroform extraction (Merck, USA; CAT. #1024451000). RNA concentration and quality were assessed using a NanoDrop ND-1000 spectrophotometer (Thermo Fisher Scientific). Reverse transcription was performed using a minimum of 500 ng of total RNA with the PrimeScript RT Master Mix kit (Clontech-Takara, Japón; CAT #RR036A). Gene expression analyses were carried out by quantitative PCR using either TaqMan probes or SYBR Green chemistry, in both cases using *GAPDH* expression as an endogenous control. TaqMan-based qRT-PCR (Thermo Fisher Scientific, USA; CAT #4331182) was used to quantify relative expression levels of *MAFG* (Hs01034678_g1) and *GAPDH* (Hs03929097_g1). Reactions were prepared using Solis Biodyne Master Mi× 10 (Genycell, Spain; CAT #08-14100001) with 2 µL of cDNA per reaction, and run on a StepOnePlus™ Real-Time PCR System (Applied Biosystems) under the following conditions: 1 cycle of 15 min at 95 °C, followed by 40 cycles of 15 s at 95 °C and 1 min at 60 °C. Genes analyzed using SYBR Green chemistry are listed in **Supplementary Table 5**, together with the specific primers designed for optimal amplification. Reactions were performed using the QuantaBio Fast PCR Kit (QuantaBio, USA; CAT #733-1387) with the following cycling conditions: 30 s at 95 °C, followed by 40 cycles of 5 s at 95 °C and 30 s at the corresponding annealing temperature (**Supplementary Table 5**). For validation of miR-4456 expression, the small RNA fraction was isolated using the miRNeasy Mini Kit (Qiagen, Germany; CAT #1038703). A total of 10 ng of small RNA was reverse-transcribed using the TaqMan™ Advanced miRNA cDNA Synthesis Kit (Thermo Fisher Scientific, USA; CAT #A28007). miR-4456 expression was quantified by qPCR using a specific TaqMan probe (Hs04274557_s1).

### Immunoblotting

Cells were washed and scraped in PBS, centrifuged, and the pellet was lysed using RIPA buffer containing protease and phosphatase inhibitor cocktail (Thermo Fisher Scientific, USA; Cat. #78440). Protein concentration was determined using a BSA standard curve and the Bradford assay (Bio-Rad, USA; CAT #5000006EDU). 30 μg of total protein were subjected to SDS-PAGE using polyacrylamide gels with a 4–15% gradient, and subsequently transferred onto Immobilon P nitrocellulose membranes (Thermo Fisher Scientific, USA; CAT #88018). Membranes were incubated overnight with primary antibodies against MAFG (Novus Biologicals, USA; CAT #nbp2-15019; 1:1000 dilution) and β-Actin (Invitrogen, Cat. #AM4302). Detection was performed using horseradish peroxidase (HRP)-conjugated secondary antibodies (1:5,000–1:10,000) and enhanced chemiluminescence (ECL) using the Pierce ECL2 Western Blotting Substrate (Thermo Fisher Scientific, USA; CAT #32209). All antibodies were diluted in 5% non-fat milk in TBS-T. Membranes were visualized using the MicroChemi 4.2 imaging system (Bioimaging Systems).

### Aptahistochemistry

MAFG protein expression was assessed by aptahistochemistry using AptMAFG3F and AptMAFG6F MAFG-specific aptamers^6^. FFPE tumor tissue sections from lung cancer patients were incubated at 60 °C for 15 min. Deparaffinization was carried out by two washes in xylene for 10 min each, followed by rehydration through graded ethanol concentrations (100%, 90%, 80%, and 70%) and distilled water for 5 min per step. Antigen retrieval was performed using heat-induced epitope retrieval in a pressure cooker for 2 min in 10 mM citrate buffer (pH 6.5). Endogenous peroxidase activity was blocked with 0.3% H_2_O_2_. Primary binding was performed overnight at room temperature using 10 pmol/mL digoxigenin-conjugated aptamers. Secondary binding was carried out with anti-digoxigenin peroxidase-conjugated antibody (Roche; 1:200 dilution in TBS) for 45 min. Signal development was performed using a DAB immunoperoxidase kit (Master Diagnostica) according to the manufacturer’s instructions. Sections were counterstained with hematoxylin.

### In silico database analyses

In silico analyses were conducted using publicly available, community-validated databases and online analytical tools. Clinical and molecular data were retrieved from The Cancer Genome Atlas (TCGA) for patients diagnosed with lung adenocarcinoma (LUAD) and lung squamous cell carcinoma (LUSC). DNA methylation data derived from Illumina HumanMethylation450 (HM450k) arrays were obtained^52^, including 460 LUAD samples and 370 LUSC samples. These data were used to evaluate methylation levels of MAFG and LIF. The UCSC Xena Browser (https://xenabrowser.net/)^53^ was used to visualize and select specific genomic regions of interest. Quantitative molecular variables, together with clinical time-to-event variables including progression-free survival (PFS), disease-specific survival (DSS), and overall survival (OS), were used to perform Kaplan–Meier survival analyses, applying the mean value of methylation or gene expression as the cut-off for patient stratification. Lists of the top 1,000 genes showing the strongest positive and negative correlations with MAFG and NFE2L2 expression were obtained for both histological subtypes using cBioPortal (https://www.cbioportal.org/)^54–56^. Gene Ontology analyses were performed and visualized using ShinyGO v0.85 (http://bioinformatics.sdstate.edu/go/)^57^ interrogating the MSigDB (Molecular Signatures Database)^58,59^

### Transfection assays

MAFG expression was modulated in cell lines using small interfering RNAs (siRNAs) (Thermo Fisher Scientific, USA; CAT #4392420) Briefly, 250,000 H23R cells were seeded in 6-well plates, and 24 h later transfected with siRNAs using the JetPRIME® transfection reagent (PolyPlus, France; CAT #101000046) according to the manufacturer’s instructions. Culture medium was replaced 24 h post-transfection. At 48 h post-transfection, cells collected to isolate DNA and RNA to confirm effective modulation of MAFG expression.

### In silico identification of MAFG-binding motifs

The intron-1 regulatory region of MAFG was analyzed for the presence of Maf Recognition Element (MARE) motifs using the FIMO algorithm from the MEME Suite (version 5.5.9). A 1,804-bp genomic fragment corresponding to the CpG-rich region within intron 1 (hg19: chr17:79,880,864–79,882,667) was extracted from the UCSC Genome Browser in FASTA format and used as input. Motif scanning was performed using the canonical MARE consensus sequence (TGCTGAGTCAGCA) and the corresponding position weight matrix (PWM) provided in MEME format. FIMO was run with default parameters, scanning both strands and applying a significance threshold of p < 1×10^−4^. Background nucleotide frequencies were estimated using the built-in non-redundant database model (--nrdb--). For each predicted motif occurrence, FIMO reported the genomic position, strand orientation, matched sequence, motif score, p-value, and false discovery rate (q-value). Genomic coordinates of motif hits were calculated by mapping the motif start and end positions to the hg19 reference genome based on the sequence header coordinates. Motif occurrences with q < 0.05 were considered significant. Visualization and schematic representation of motif locations relative to the CpG cluster were generated manually based on the FIMO output.

### Statistical analysis

Parametric or non-parametric statistical tests were used, as appropriate, to determine statistical significance across all experiments in this study. Data from expression assays are presented as mean ± standard deviation (SD) of three technical replicates from at least two independent experiments. DNA methylation data are presented as the mean of two technical replicates from at least two independent experiments. Welch’s two-tailed t-test was applied to identify statistically significant differences between two groups of quantitative data. Pearson or Spearman correlation analyses were performed to assess associations between quantitative variables, depending on data distribution. The Shapiro–Wilk test was used to evaluate data normality. For Kaplan–Meier survival analyses, statistical significance was assessed using the Log-Rank test. A p value < 0.05 was considered statistically significant in all analyses. In gene ontology (GO) enrichment analyses, a false discovery rate (FDR) < 0.1 was applied for the GO1 and GO3 datasets due to its limited sample size, whereas an FDR < 0.05 was used for all other enrichment analyses. All gene sets associated with transcriptional signatures linked to MAFG and NFE2L2 met a q-value (adjusted *p*-value) < 0.05. Statistical computations were performed using SPSS software version 20 (IBM, USA) and GraphPad Prism version 9.0.0 (GraphPad Software, USA).

## DECLARATIONS

### Ethics approval and consent to participate

All samples were processed following the standard operating procedures with the appropriate approval of the Human Research Ethics Committees, including informed consent within the context of research (HULP: PI-5063).

### Availability of data and materials

The datasets generated and/or analyzed during the current study are available in the GEO repository (number pending).

### Declaration of generative AI and AI-assisted technologies in the manuscript preparation process

During the preparation of this work the authors used Copilot to enhance readability and improve the overall flow of the manuscript. After using this tool, the authors reviewed and edited the content as needed and take full responsibility for the content of the published article.

### Authors contributions

Conceptualization, AGG, IIC, OV. Methodology, AGG, CRA, AAC, RMV, OP, MB, LAR, SS, RTR, SRP. Formal analysis: AGG, CRA, OV. Investigation, AGG, CRA, AAC, RMV, OP, MB, LAR, IER, SS, RTR, SRP, ASP, VMG, JDC, IIC, OV. Resources IER, JDC, IIC, OV. Writing original draft, AGG, IIC, OV. Writing, review and editing, AGG, CRA, AAC, RMV, OP, MB, LAR, SS, IER, RTR, SRP, ASP, VMG, JDC, IIC, OV. Supervision, OV, IIC. Project administration, OVP, IIC. Funding acquisition, JDC, IIC, OV.

### Authorship

We declare that all the authors of this study have directly participated in the planning, execution, or analysis of the study, and all the authors have read and approved the final version submitted, adhering to the guidelines of the ICMJE.

## Acknowledgments

The authors thank HULP-IdiAPZ Biobank for sample processing and the cell culture platform at IdiPAZ. We also thank the Biostatistics Platform at IdIPAZ, especially Itsaso Losantos-García for her dedication. We are also grateful for the financial support.

