## Supplementary Figures for "Epigenetic silencing of MAFG is a potential prognosis biomarker in lung adenocarcinomas"

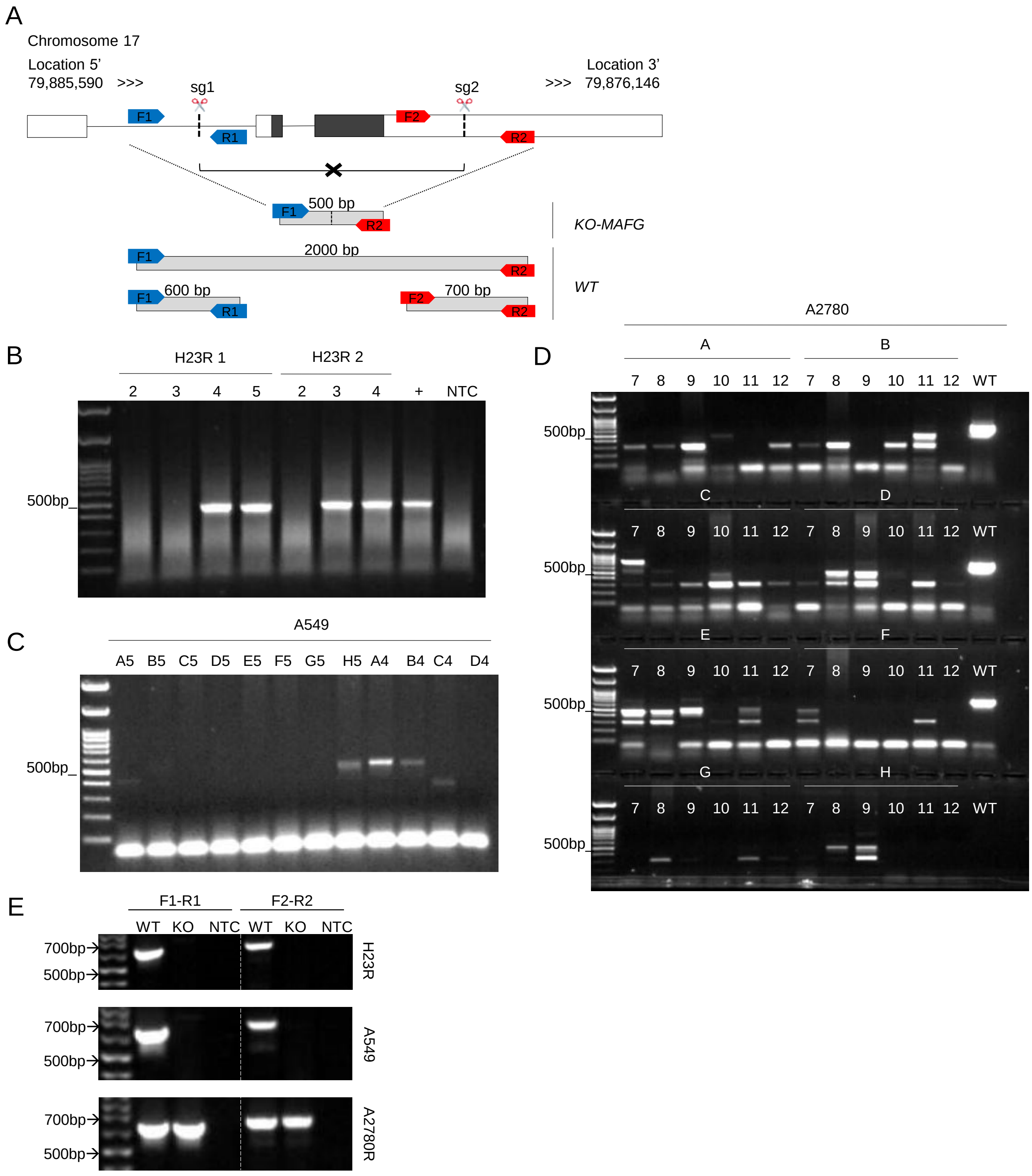


**Supplementary Figure 1. PCR screening of CRISPR/Cas9 MAFG-KO clones**. **(A)** Schematic representation of the MAFG genomic locus and CRISPR/Cas9 strategy. Two single-guide RNAs (sg1 and sg2) flanking the MAFG coding sequence were designed, together with primer pairs F1–

R1 (sg1 site), F2–R2 (sg2 site), and F1–R2 to discriminate wild-type (Ctrl) and MAFG knockout (KO) alleles. **(B-D)** Semi-quantitative PCR amplification using primer pairs F1–R2 in genomic DNA from H23R (B), A549 (C) and A2780R (D) cell lines after the CRISPR/Cas9 process. **(E)** Semi-quantitative PCR amplification using primer pairs F1–R1 and F2–R2 in genomic DNA from Ctrl and MAFG-KO clones derived from H23R, A2780R and A549 cell lines.

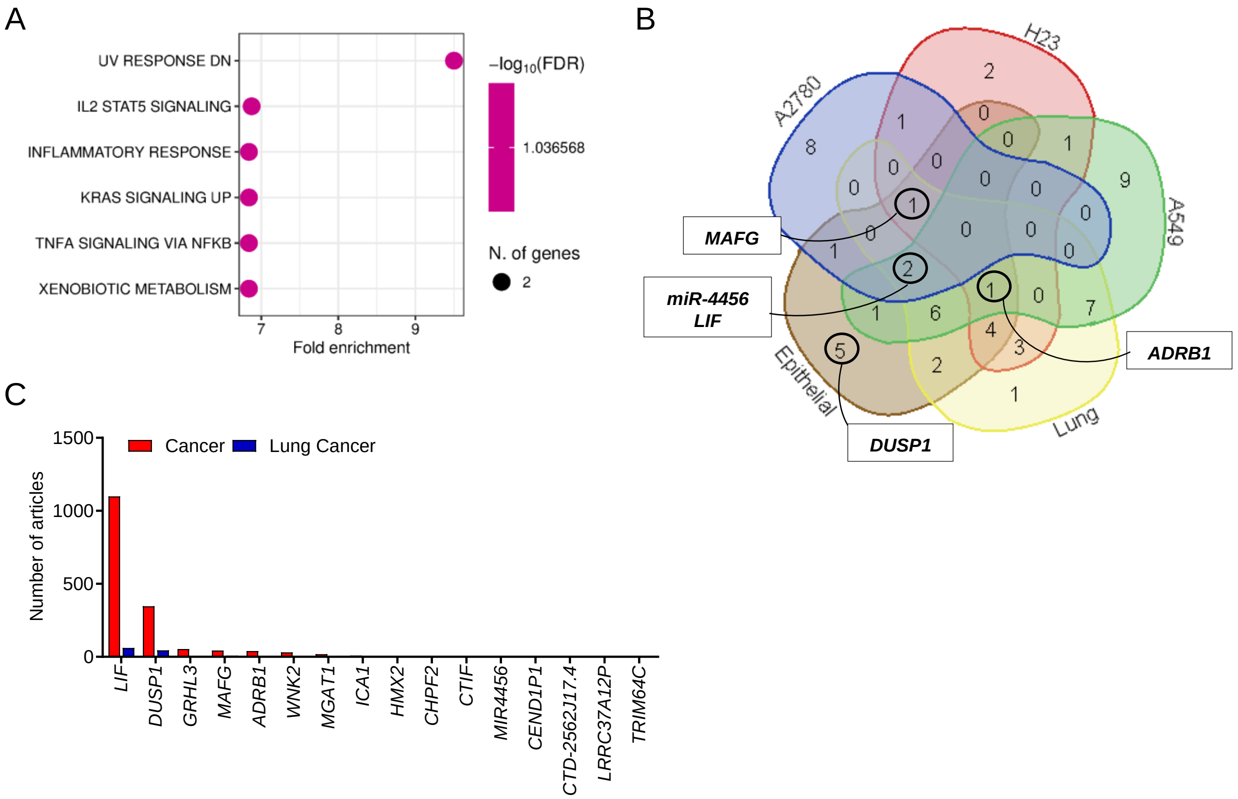


**Supplementary Figure 2. Molecular alterations derived from MAFG-KO**. **(A)** Pathway enrichment analysis of the 56 differentially methylated genes identified by SeqCap after MAFG deletion was performed using ShinyGO v0.85 (Ge SX, Jung D & Yao R, Bioinformatics 36:2628–2629, 2020) with the Hallmark gene sets from MSigDB and an FDR cutoff of 0.1. **(B,C)** Venn diagram showing the distribution of candidate genes identified across individual cell lines (H23R, A2780R, A549), combined lung cancer analysis ("LUNG"), and combined epithelial analysis ("Epithelial"). **(D)** Number of articles for each selected gene associated to "Cancer" and "Lung Cancer" according to PUBMED (Last search on November 2024).

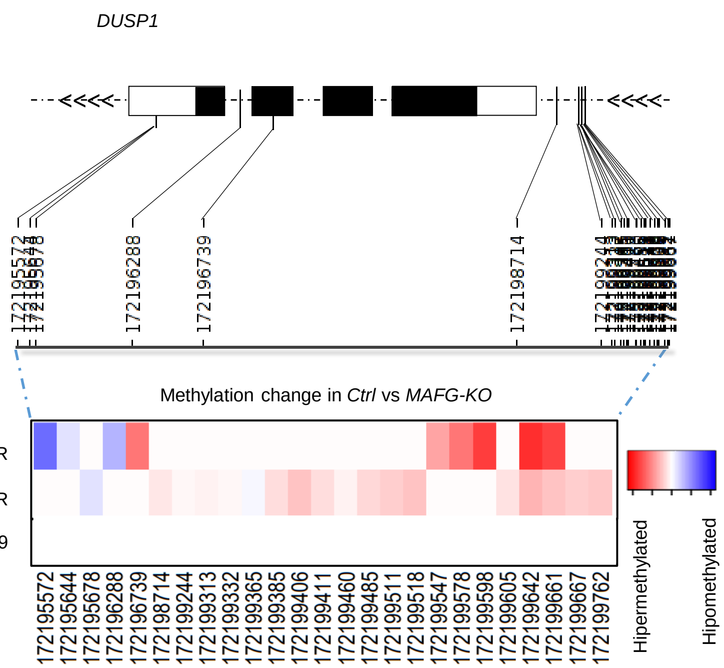


**Supplementary Figure 3. Methylation changes in DUSP1 after MAFG deletion**. Methylation changes observed in SeqCap of CpG sites located in the promoter region of DUSP1 MAFG-KO vs Ctrl cells.

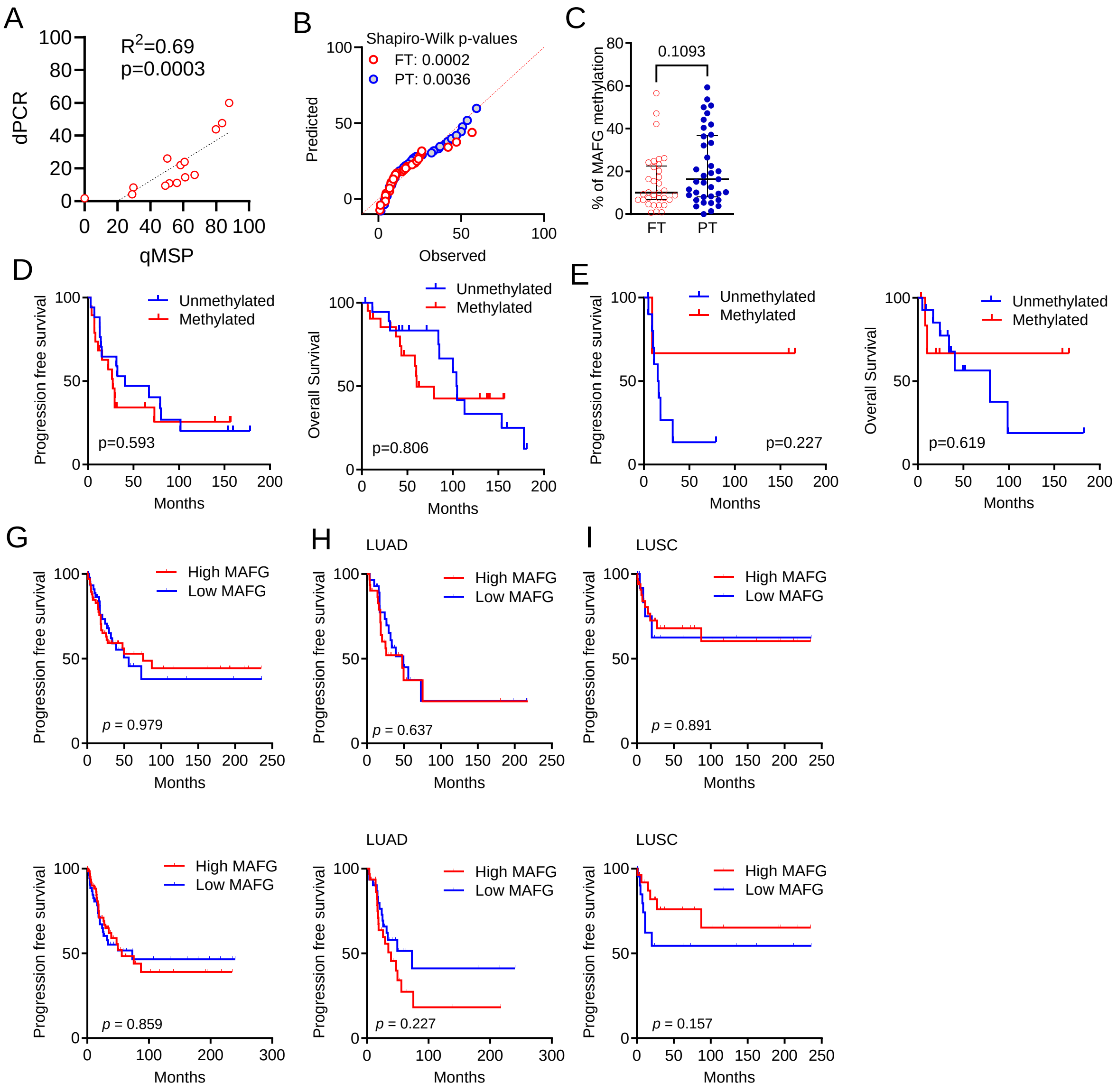


**Supplementary Figure 4. Clinical relevance of MAFG in NSCLC patients. (A)** Correlation between MAFG methylation levels measured by qMSP and digital PCR (dPCR) in fresh tumor samples from NSCLC patients. **(B,C)** Normality (B) and mean (C) analyses of MAFG methylation levels in fresh-frozen (FT, n=33) and FFPE (PT, n=37) NSCLC samples. **(D,E)** Kaplan–Meier analyses of progression-free survival (PFS) and overall survival (OS) in an internal NSCLC cohort, stratified by MAFG methylation levels, for LUAD (D) and LUSC (E). **(F)** Kaplan–Meier analysis of progression free survival (PFS) in 127 early-stage NSCLC patients based on MAFG levels detected using AptMAFG3F (top) or AptMAFG6F (bottom). **(G-I)** Subgroup Kaplan–Meier analyses according to histological subtype, showing PFS in LUAD (H,J) and LUSC (I,K) patients stratified by MAFG expression detected with AptMAFG3F (H,I) or AptMAFG6F (J,K). Median expression values were used as cut-off thresholds. Log-rank (Mantel-Cox) test was used to determine differences in survival between groups.

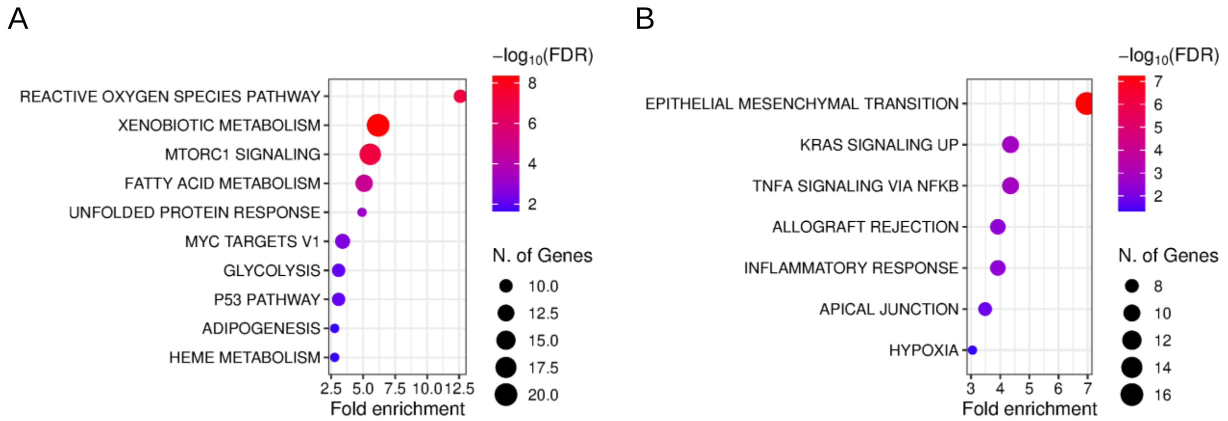


**Supplementary Figure 5. Molecular pathways associated to the transcriptional signature of MAFG and NFE2L2 in LUSC (A,B)** Molecular pathways enriched among genes shared between the positive (A) and negative (B) transcriptional signatures of *MAFG* and *NFE2L2* in lung squamous cell carcinoma (LUSC). Pathway enrichment analysis was performed using ShinyGO v0.85 (Ge SX, Jung D & Yao R, Bioinformatics 36:2628–2629, 2020) with the Hallmark gene sets from MSigDB and an FDR cutoff of 0.05.
